# Tbr2-expressing retinal ganglion cells are ipRGCs

**DOI:** 10.1101/2020.06.17.153551

**Authors:** Chai-An Mao, Ching-Kang Chen, Takae Kiyama, Nicole Weber, Christopher M. Whitaker, Ping Pan, Tudor C. Badea, Stephen C. Massey

**Affiliations:** Ruiz Department of Ophthalmology and Visual Science, McGovern Medical School at The University of Texas Health Science Center at Houston (UTHealth), Houston, TX 77030, USA; Department of Ophthalmology, Baylor College of Medicine, Houston, TX, 77030, USA; National Eye Institute, National Institutes of Health, Bethesda, Maryland 20892, USA

## Abstract

The mammalian retina contains more than 40 retinal ganglion cell (RGC) subtypes based on their unique morphologies, functions, and molecular profiles. Among them, intrinsically photosensitive RGCs (ipRGCs) are the first specified RGC type that emerged from a common pool of retinal progenitor cells. Previous work has shown that T-box transcription factor *T-brain 2* (*Tbr2*) is essential for the formation and maintenance of ipRGCs, and Tbr2-expressing RGCs activate *Opn4* expression upon native ipRGC loss, suggesting that Tbr2^+^ RGCs can serve as a reservoir for ipRGCs. However, the identity of Tbr2^+^ RGCs has not been fully vetted, and the developmental and molecular mechanisms underlying the formation of native and reservoir ipRGCs remain unclear. Here, we showed that Tbr2-expressing retinal neurons include RGCs and GABAergic displaced amacrine cells (dACs). Using genetic sparse labeling, we demonstrated that the majority of Tbr2^+^ RGCs are intrinsically photosensitive and morphologically indistinguishable from known ipRGC types and have identical retinofugal projections. Additionally, we found a minor fraction of Pou4f1-expressing Tbr2^+^ RGCs marks a unique OFF RGC subtype. Most of the Tbr2^+^ RGCs can be ablated by anti-melanopsin-SAP toxin in adult retinas, supporting that Tbr2^+^ RGCs contain reservoir ipRGCs that express melanopsin at varying levels. When *Tbr2* is deleted in adult retinas, *Opn4* expression is diminished followed by the death of *Tbr2*-deficient cells, suggesting that Tbr2 is essential for both *Opn4* expression and ipRGC survival. Finally, Tbr2 extensively occupies multiple *T*-elements in the *Opn4* locus, indicating a direct regulatory role for Tbr2 on *Opn4* transcription.

**Significance statement:** Melanopsin/Opn4-expressing intrinsically photosensitive retinal ganglion cells (ipRGCs) play fundamental roles in non-image forming vision. Previously we identified *Tbr2* as the key transcription regulator for the development and maintenance of ipRGCs. To reveal the full identity of Tbr2-expressing retinal neurons and how Tbr2 acts, we generated a novel mouse line to genetically label and study Tbr2-expressing cells. Our in-depth characterizations firmly established that most Tbr2^+^ RGCs are indeed ipRGCs and that Tbr2 regulates *Opn4* transcription, thus place Tbr2-Opn4 transcription regulatory hierarchy as the primary component in the development and maintenance of the non-image forming visual system.

## Introduction

Intrinsically photosensitive RGCs (ipRGCs) are the first specified RGC types marked by the expression of *opsin 4* (*Opn4*, aka melanopsin) in developing retina starting at embryonic day 14 (E14) (McNeill et al., 2011, Mao et al., 2014). The primary function of ipRGCs is to enable non-image-forming visual functions, such as pupillary light reflex (PLR) and circadian photo-entrainment (Lazzerini Ospri et al., 2017). They also play a role in light-dependent exacerbation of migraines (Noseda et al., 2010), sleep deprivation (Tsai et al., 2009, Altimus et al., 2008), and mood swing (LeGates et al., 2014).

Previously we and others have identified *T-brain 2* (*Tbr2* or *Eomes*) as a key regulator in ipRGC formation (Mao et al., 2008, Sweeney et al., 2014, Mao et al., 2014). Many Tbr2-expressing RGCs can activate *Opn4* expression upon native ipRGC loss, suggesting that a reservoir for ipRGCs exist among Tbr2^+^ RGCs (Mao et al., 2014). However, several questions remain unanswered regarding the identity of Tbr2^+^ RGCs and the role Tbr2 plays in ipRGC development. First, an estimated 17.8% of all RGCs were thought to be Tbr2^+^ RGCs, yet only ~40% of Tbr2^+^ RGCs appeared as ipRGC reservoir (Mao et al., 2014). The identities of the other 60% of Tbr2^+^ RGCs were not determined. Second, it has been shown that all supra-chiasmatic nucleus (SCN)-afferent RGCs expressed melanopsin (Baver et al., 2008), contradicting our observation that many Tbr2^+^ reservoir ipRGCs do not express melanopsin or they do but at levels below detection by conventional immunostaining. Third, our conclusion that *Tbr2* is essential for maintaining ipRGC survival was drawn from the near-complete loss of melanopsin-expressing cells in *Opn4^Cre^:Tbr2^f/f^* retinas. Could this phenotype be interpreted as a requirement for *Tbr2* in ipRGC development as *Opn4* is also expressed in developing RGCs at embryonic stages? Finally, what are the roles of *Tbr2* in regulating *Opn4* expression in ipRGC reservoir in adult retinas? In this report, we shall answer these questions.

To determine the identity of Tbr2^+^ RGCs and to study Tbr2’s functions in ipRGC development, we generated a novel knock-in mouse line with the engineered *Tbr2^TauGFP-IRESCreERT2^* allele shown in Fig. 1A. Using this mouse line in conjunction with *Rosa^iAP^, Pou4f1^CKOAP^*, and *Ai9* reporter lines, we conducted sparse labeling genetically to investigate *Tbr2*-expressing cells. We found that the majority of Tbr2^+^ RGCs are morphologically indistinguishable from known ipRGCs, including the M1-M6 subtypes. Evidently, these Tbr2^+^ RGCs project to the same regions as known ipRGCs, including SCN, ventral lateral geniculate nuclei (vLGN), inter-geniculate leaflet (IGL), and olivary pretectal nuclei (OPN). We also found a small fraction of Pou4f1-expressing Tbr2^+^ RGCs, which we morphologically identified as a unique bushy OFF RGC subtype with dendritic stratification distinctively below those of conventional M1 type. In addition to RGCs which constitute ~46% of Tbr2^+^ neurons, we surprisingly found that ~54% of Tbr2^+^ cells are actually GABAergic displaced amacrine cells (dACs). This new data explains a mystery in our previous report where only ~40% of Tbr2^+^ cells were depleted in *Opn4^Cre^:Tbr2^f/f^* retinas (Mao et al., 2014). Moreover, by molecular surgery in adult retinas, a significant fraction (approximately 40%) of Tbr2^+^ cells can be ablated by SAP-melanopsin, substantiating the notion that the majority of Tbr2^+^ RGCs express melanopsin and are indeed ipRGCs. By loss-of-function studies, we extended our previous finding that *Tbr2* is indeed essential for the survival of adult Tbr2^+^ RGCs. More importantly, we found that Opn4 expression is significantly down-regulated prior to the death of *Tbr2*-deleted RGCs in adult retinas. Finally, we showed that Tbr2 binds to multiple *T*-elements in the *Opn4* locus, thus demonstrating a direct role of Tbr2 in *Opn4* expression.

**Figure 1.**
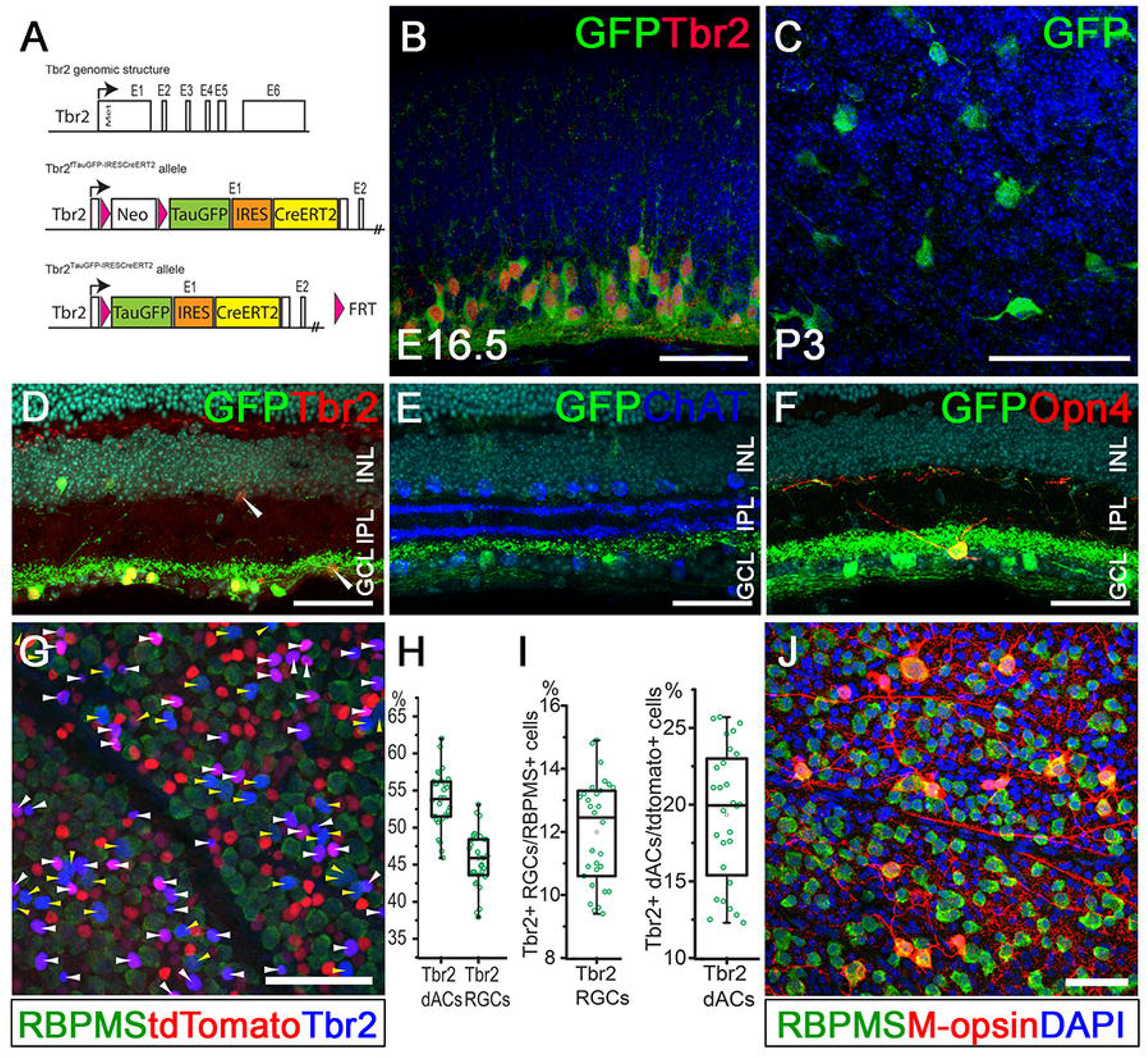
Tracing Tbr2-expressing retinal neurons. **(A)** Genomic structure of *Tbr2* locus and the engineered allele in the *Tbr2^TauGFP-IRESCreERT2^* mouse. **(B)** Double immunofluorescent staining showing Tbr2 expression (red) in Tbr2-driven GFP^+^ cells (green) in an E16.5 *Tbr2^TauGFP^* retinal section. **(C)** Representative immunofluorescent image showing Tbr2-driven GFP^+^ cells (green) in a P3 *Tbr2^TauGFP^* retinal flatmount. **(D)** Immunofluorescent image showing Tbr2 expression (red) in Tbr2-driven GFP^+^ cells (green) in a *Tbr2^TauGFP^* retinal section. **(E)** Fluorescent image of a *Tbr2^TauGFP^* retinal section showing dense Tbr2-driven GFP signal (green) in ON sub-laminae below the cholinergic ChAT bands (blue) in inner plexiform layer (IPL). **(F)** Double immunofluorescent staining on a *Tbr2^TauGFP^* retinal section showing melanopsin expression (red) in Tbr2-driven GFP^+^ cells (green). **(G)** Co-immunofluorescent staining on a *Slc32a1^Cre^:Ai9* retinal flatmount showing Tbr2 expression (blue) with RBPMS^+^ RGCs (yellow arrowheads) or tdTomato^+^ dACs (white arrowheads). **(H)** Relative abundance of Tbr2^+^ RGCs and dACs described in G. **(I)** Relative abundance of Tbr2^+^ RGCs within the entire RGC population (left) and Tbr2^+^ dACs within the entire dAC population. **(J)** Co-immunofluorescent staining on a *WT* retinal flatmount showing melanopsin expression (red) with RBPMS^+^ RGCs (green). ChAT: cholinergic acetyltransferase. GCL: ganglion cell layer. INL: inner nuclear layer. IPL: inner plexiform layer. Scale bars: 50 μm (B-J).

## Materials and Methods

### Animals

*Tbr2^TauGFP-IRESCreERT2^* mice (described in Fig. 1A) were generated by inserting a Frt-PGKNeopA-Frt cassette and a synthetic *TauGFP-IRESCreERT2* DNA fragment in the first exon behind the translation initiation codon of *Tbr2*. Gene targeting was conducted as described previously (Mao et al., 2008). Two targeted embryonic stem (ES) cell lines were used to generate *Tbr2^fTauGFPIRESCreERT2^* mice. The *Tbr2^fTauGFPIRESCreERT2^* mice were subsequently bred to a *R26-FLPeR* line to delete the *Frt-flanked PGKNeopA* cassette to generate the *Tbr2^TauGFPIRESCreERT2^* mice. PCR primers Tbr2f1 and Taur1 were used for genotyping the *Tbr2^TauGFPIRESCreERT2^* allele (Table 1). *Tbr2^TauGFPIRESCreERT2^* mice were bred to *Rosa^iAP^* or *R26-tdTomato* (*Ai9*) mice to produce double-transgenic mice for sparse labeling, dye filling and patch clamp recording. Tamoxifen (1-5 consecutive daily injections of 100 μg/g body weight; Sigma, St. Louis, MO) was injected intraperitoneally for Cre induction. Alternatively, 1 μl of 4-hydroxytamoxifen (0.1 mg/ml; Sigma) was injected into the vitreous space of the right eye to activate cells locally. The generation and genotyping of *Rosa^iAP^, Pou4f1^CKOAP^, Ai9, R26^FLPeR^, Slc32a1^Cre^, Opn4^Cre^*, and *Opn4^TauLacZ^* mice were described previously (Badea et al., 2009a, Madisen et al., 2010, Farley et al., 2000, Vong et al., 2011, Ecker et al., 2010, Hattar et al., 2002). All mice were maintained on C57BL6/129 mixed backgrounds. Mouse lines of either sex at various ages between P21 to 6 months were used. Pre-weaned animals were housed with their mother; weaned animals were housed in groups of no more than 5 in individually ventilated cages. All animal procedures followed the US Public Health Service Policy on Humane Care and Use of Laboratory Animals and were approved by the Animal Welfare Committee at The University of Texas Health Science Center at Houston and Institutional Animal Care and Use Committee at Baylor College of Medicine.

**TABLE 1.**
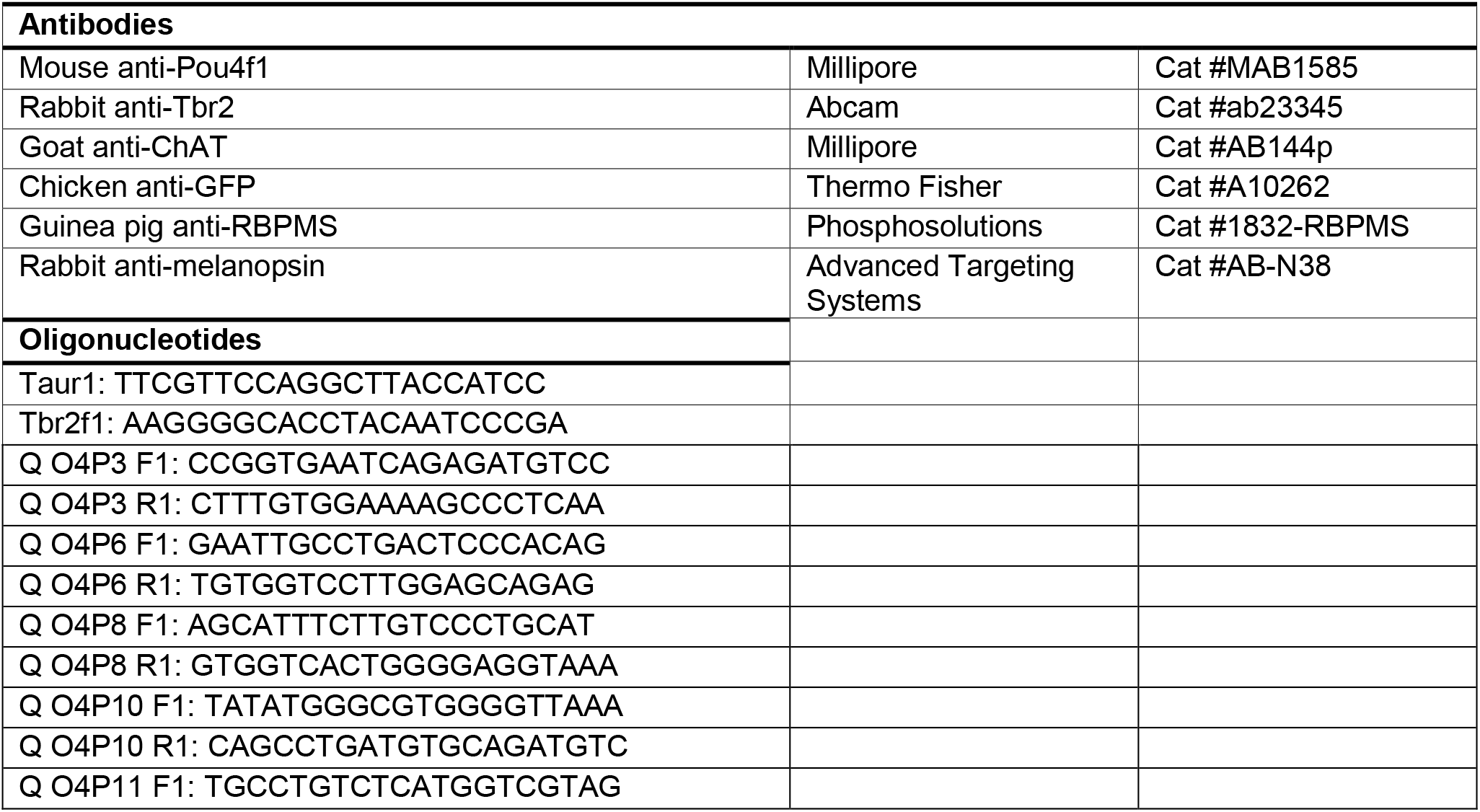
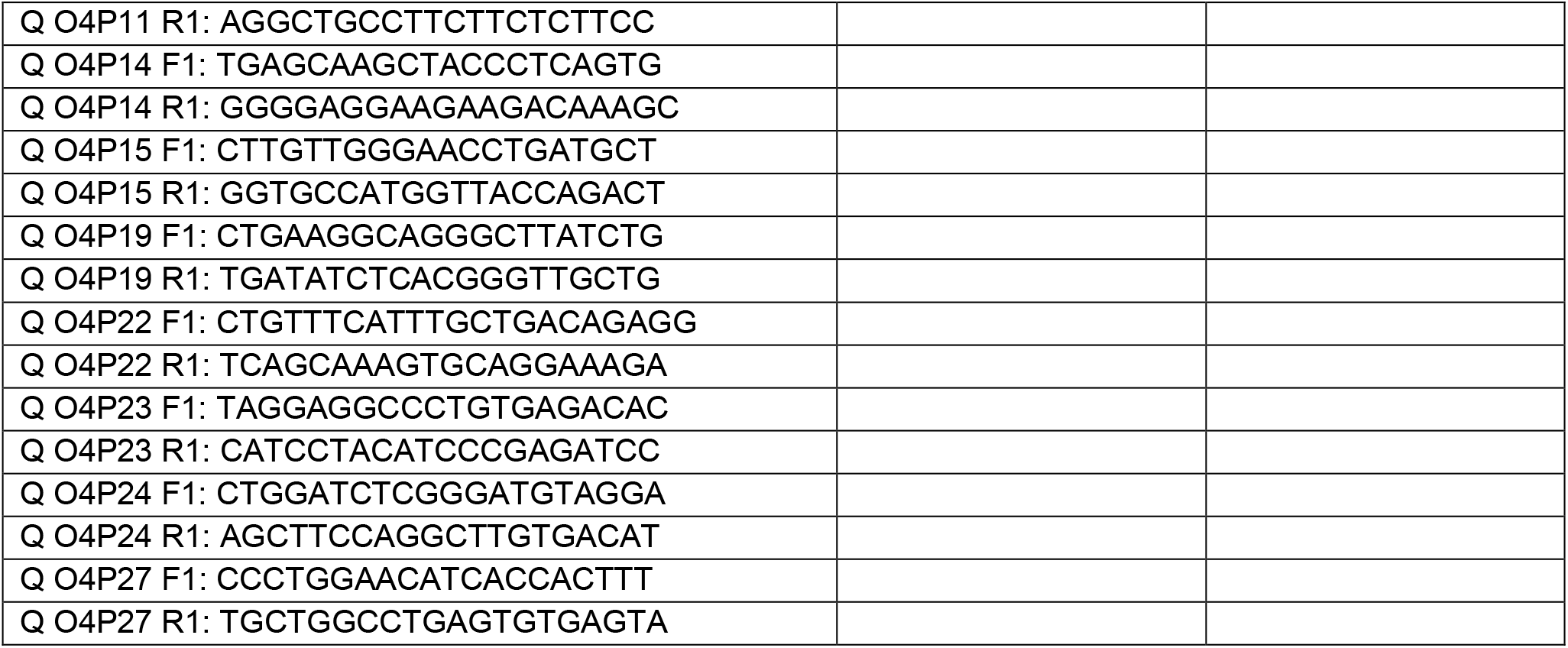
Antibodies and Oligonucleotides

### Immunohistochemical analysis

Retinal sections or flat-mounted retinas were fixed with 4% paraformaldehyde (PFA; Electron Microscopy Sciences, Hatfield, PA), then incubated with the primary antibodies listed in Table 1. Alexa Fluor-conjugated secondary antibodies were obtained from Life Technologies, Carlsbad, CA (1:600 dilution). DAPI (2.5 μg/ml) was used to stain the nuclei. Images were acquired on a Zeiss 780 confocal laser scanning microscope (Carl Zeiss, Thornwood, NY) and exported as TIFF files into Adobe Photoshop (Adobe Systems, San Jose, CA). Cell counting was conducted using the cell counter plugin of NIH ImageJ.

### Alkaline phosphatase (AP) staining

Tamoxifen-treated *Tbr2^TauGFP/+^:Rosa^iAP/+^* mice were used for alkaline phosphatase (AP) staining, which was conducted as previously described with minor modifications (Badea et al., 2003, Jamal et al., 2020). Whole eyeballs were fixed with 10% neutrally buffered formalin for 5 min. The retinas were removed and flat mounted on a piece of nitrocellulose membrane, post-fixed for 10 min at room temperature, washed twice in phosphate-buffered saline (PBS), and heated in PBS for 30 min in 65°C water bath to inactivate endogenous AP activity. AP staining was performed in 0.1 M Tris (pH 9.5), 0.1 M NaCl, 50 mM MgCl_2_, 0.34 g/ml p-nitroblue tetrazolium chloride (NBT), and 0.175 g/ml 5-bromo-4-chloro-3-indolyl phosphate (BCIP) solution for 24-48 hours at room temperature with gentle shaking. After staining, retinas were washed 3 times for 5 min in PBS, post-fixed in PBS with 4% PFA briefly, dehydrated through an ethanol series, and then cleared with 2:1 benzyl benzoate/benzyl alcohol.

Montages of the whole retina were acquired on a Zeiss Axio Imager 2 microscope equipped with a motorized xyz drive (Carl Zeiss, White Plains, NY). Individual RGCs were imaged under transmitted light, and z-stacks were collected with a 20X objective (3.1 pixels/μm) at 0.58 μm vertical (z-dimension) intervals. For each RGC, the areas of dendritic arbors and somata, the branching, and the stratification depth within the inner plexiform layer (IPL) were analyzed using Zen lite (Carl Zeiss). Diameters of dendritic arbors and cell bodies were measured for each cell by calculating the diameter of a circle having the same area as a convex polygon, minimally enclosing the dendritic or somatic profile. Individual examples of distinct morphological types for M2, M4, and M5-like cells were traced using Adobe Photoshop or the NeuronJ plugin of ImageJ (Meijering et al., 2004).

### Tracing RGC axon targets

Two μl of cholera toxin subunit B (CTB) conjugated with Alexa Fluor-555 (1 mg/ml) was injected into the vitreous of the right eye of *Tbr2^TauGFP/+^* mice using a 35-gauge NanoFil system (World Precision Instruments, Sarasota, FL). Two days after CTB injection, animals were anesthetized and perfused with 4% PFA. For inspecting *Tbr2^CreERT2^*-driven *Rosa^iAP^* expression in brains, tamoxifen-treated mice were perfused with 4% cold PFA, and then the whole brains were dissected and post-fixed with 4% PFA. Cryopreserved brains were sectioned into consecutive 100 μm coronal sections for additional processing using either immuno-staining with GFP antibody on *Tbr2^TauGFP/+^* brain sections or AP staining on *Tbr2^TauGFP/+^:Rosa^iAP/+^* brain sections.

### Saporin (SAP)-induced molecular surgery

One μl of anti-melanopsin conjugated with saporin (Melanopsin-SAP, 400 μg/ml, Advance Targeting System, San Diego, CA) was injected into the vitreous of the right eye of *Tbr2^TauGFP/+^* mice using a 33-gauge NanoFil system (World Precision Instruments, Sarasota, FL). The left eye was injected with one μl of rabbit IgG-SAP as a negative control. One month later, the same procedure was repeated. Two months after the first injection, retinas were isolated for immunofluorescent analysis for GFP and melanopsin expression.

### Dye filling

Dye filling was conducted under room light as previously described with minor modifications (Mao et al., 2014). Briefly, retinas were flat-mounted on black membrane filter paper, and perfused with oxygenated Ames’ medium (US Biological, Salem, MA) in a recording chamber. The glass microelectrodes were tip-filled with a mixture of 0.5% Lucifer yellow-CH and 3.5% NBT and then backfilled with 3 M LiCl. The tdTomato^+^ RGCs were viewed under an epifluorescence microscope with a 40X water immersion objective and randomly targeted. Cell penetration was verified within the first minute of iontophoresis of Lucifer yellow with negative current (−0.5 nA at 3 Hz). NBT was then injected into cells with a positive current (+0.5 nA at 3Hz) for 5-10 min. Tissue was fixed with 4% PFA in 0.1 M PBS for 1 hr at room temperature. Filled cells were visualized by incubating with Alexa Fluor 488-conjugated streptavidin (1:1000 dilution) at 4°C overnight. The injected retinas were stained by an anti-GFP antibody to verify the targeted RGCs and an anti-ChAT antibody to determine the depth of dendritic stratification of injected cells.

### Intrinsic membrane property and photosensitivity

Whole cell patch clamp recording was used as described in Kiyama et al. (2019) to test intrinsic photosensitivity of Tbr2^+^ cells. Retinas from tamoxifen-treated *Tbr2^CreERT2/+^:Ai9* mice were flat-mounted with ganglion cell side up under dim red light and perfused (3 ml/minute) with a warm (~34°C) carbogenated mammalian Ringer solution containing (in mM) 120 NaCl, 5 KCl, 25 NaHCO_3_, 0.8 Na_2_HPO_4_, 0.1 NaH_2_PO_4_, 2 CaCl_2_, 1 MgSO_4_, 10 D-glucose, 0.01 L-AP4, 0.01 AP5 and 0.04 NBQX) on an Olympus BX51WI fixed-stage microscope (Olympus USA, Central Valley, PA, USA). Retinal neurons were visualized under differential interference contrast with infrared illumination (> 900 nm) and Tbr2^+^ cells marked by tdTomato expression were identified by 50 msec light pulses (35000 R*/rod/sec) delivered and imaged through the mCherry filter set (Olympus USA). The cumulative exposure time was less than 1 second in order to preserve light sensitivity. Once a cell was selected, a 20-minute wait time was exercised to allow dark adaptation before recording. Patch pipettes made from borosilicate glass tubes (8-12 MΩ) were filled with a potassium-based internal solution containing (in mM) 125 K-gluconate, 8 NaCl, 4 ATP-Mg, 0.5 Na-GTP, 5 EGTA, 10 HEPES and 0.2% biocytin (w/v) with pH adjusted to 7.3 using KOH. The liquid junction potential was 15 mV and not corrected. Current clamp recording was conducted using the AM2400 amplifier (A-M Systems, Sequim, WA, USA) driven by the WinWCP program provided by John Dempster (University of Strathclyde, Glasgow, UK). Signals were filtered at 5 kHz and sampled at 10 kHz. To obtain intrinsic membrane property (IMP), a small negative current was injected immediately after break-in to adjust membrane potential from rest to −64 ± 2 mV. This was then followed by 5 serial negative current injections of 600 msec in duration for cell hyperpolarization, followed by 4 serial positive current injections for depolarization. The amount of negative current required to hyperpolarize a cell to 100 ± 5 mV was empirically determined for each cell and used to adjust the amplitudes of subsequent current steps. Under these conditions, Tbr2^+^ cells exhibit characteristic ensemble IMP profiles that match cells with similar dendritic morphology and IPL stratification levels. Intrinsic photoresponse was determined by a 1-sec 470 nm light stimulation (~430 μm in diameter) at intensities of 2.2 to 5.9 log photon/μm^2^. The dendritic morphology of the most sensitive M1-like cells were recovered by *post hoc* staining for biocytin and IPL stratification level was determined in reference to two ChAT bands as described above. Dendritic structure was then traced and quantified in Neurolucida 360 and Neurolucida Explorer programs (MBF Bioscience, Boston, MA).

### Chromatin immunoprecipitation

Twenty-two wild-type (WT) P0 retinas were isolated for chromatin immunoprecipitation (ChIP) assays as described (Tsai et al., 2008) with slight modification. Briefly, retinas were cross-linked with 1% formaldehyde for 10 min, stopped by 0.125 M glycine and then homogenized in the cell lysis buffer (5mM PIPES pH 8.0, 85 mM KCl, 0.5% NP40, and proteinase inhibitors). Nuclei were collected and resuspended in the nuclei lysis buffer (50 mM Tris-HCl pH 8.1, 10 mM EDTA, 1% SDS, and proteinase inhibitors). Chromatin was sheared by a Diagenode Bioruptor Plus sonication system (Diagenode, Denville, NJ). Fragmented chromatin was precleared with 2.5 μg of normal rabbit IgG, then incubated overnight with 1 μg of rabbit anti-Tbr2 antibody or normal rabbit IgG (Santa Cruz Biotechnology, Dallas, TX). Antibody-bound chromatin complex was precipitated with salmon sperm DNA/Protein A agarose (EMD Millipore, Burlington, MA), then washed sequentially with PIRA (150 mM NaCl, 50 mM Tris-HCl pH 8.0, 0.1% SDS, 0.5% deoxycholate, 1% NP40, and 1 mM EDTA), high salt buffer (50 mM Tris-HCl pH 8.0, 500 mM NaCl, 0.1% SDS, 0.5% deoxycholate, 1% NP40, and 1 mM EDTA), LiCl wash buffer (50 mM Tris-HCl pH 8.0, 1 mM EDTA, 250 mM LiCl, 1 % NP40, and 0.5% deoxycholate) and TE for 10 min each at 4°C. Precipitated DNA was treated with RNase A and proteinase K at 37°C overnight. Cross-linking was reversed at 65°C overnight. DNA was extracted by phenol/chloroform, precipitated with ethanol, and dissolved in 30 μl of water. Three μl of DNA solution was used for one real-time quantitative PCR (qPCR) reaction. To analyze specific Tbr2-bound DNA, we performed qPCR using the CFX Connect Real-Time PCR Detection System with iTaq Universal SYBR Green Supermix (Bio-Rad, Hercules, CA). The qPCR primers are described in Table 1.

## EXPERIMENTAL DESIGN AND STATISTICAL ANALYSIS

Mouse lines (*Tbr2^TauGFP-IRESCreERT2/+^: Rosa^iAP/+^* and *Tbr2^TauGFP-IRESCreERT2/+^: Ai9*) of either sex at various ages between P20 to 6 months were used for sparse labeling, dye filling and patch clamp recording experiments. Cell counting experiments described in Fig.1 were conducted on 3 *Slc32a^Cre/+^:Ai9* mice. All data are presented as mean ± standard deviation for each genotype. For comparing the number of Tbr2^+^ RGCs after melanopsin-SAP ablation, a two-tailed one-sample Student’s *t*-test in Excel (Microsoft, Redmond, WA) was used. Results were considered significant when *P*<0.05.

## Results

### Tbr2-driven TauGFP expression recapitulates endogenous Tbr2 expression

In order to trace retinal neurons that express *Tbr2*, we have generated a new knock-in mouse line *Tbr2^TauGFP-IRESCreERT2^* (labeled as *Tbr2^TauGFP^* or *Tbr2^CreERT2^* interchangeably in the text according to its usage relevancy) in which a *TauGFP-IRESCreERT2* dual reporter cassette was inserted in-frame behind the translational start codon of *Tbr2* (Met, Fig. 1A). Immunostaining with an anti-GFP antibody on developing retinas showed that GFP could be found in E16.5 and P3 *Tbr2^TauGFP/+^* retinas (Fig. 1B, 1C). In adult *Tbr2^TauGFP/+^* retinas, GFP were clearly present in most Tbr2-expressing cells and the signal weakened when Tbr2 expression levels were barely detectable (white arrowheads in Fig. 1D). These data validated the use of this new mouse line to track endogenous *Tbr2* expression. We found in adult *Tbr2^TauGFP^* retinal sections that GFP signal was conspicuously present in the lower part of IPL below the ON cholinergic acetyltransferase (ChAT) band. A weaker but clearly noticeable signal is found in a narrow band lining the border between IPL and inner nuclear layer (INL) (Fig. 1E) similar to that of Opn4/melanopsin staining patterns. We then compared Opn4 and GFP expression on *Tbr2^TauGFP^* retinal sections and found that all detectable soma and IPL Opn4 signal coincided with GFP (Fig. 1F), further affirming the use this mouse line for tracking Tbr2 expression and studying the relationship between *Tbr2* and melanopsin-expressing ipRGCs.

Previous estimate put Tbr2-expressing cells at approximately 8.9% of all DAPI^+^ nuclei in the ganglion cell layer (GCL) (Mao et al., 2014). The conclusion that Tbr2^+^ RGCs account for 17.8% of all RGCs was based on 2 assumptions: 1) All Tbr2^+^ cells are RGCs, and 2) RGCs account for ~50% of all cells in the GCL. To test these assumptions, we co-stained GFP with RBPMS, a pan-RGC marker (Kwong et al., 2010). Surprisingly, we found that less than half of GFP^+^ cells expressed RBPMS (data not shown), suggesting that only a fraction of Tbr2^+^ cells are RGCs and that the Tbr2^+^RBPMS^-^ cells are displaced amacrine cells (dACs). To verify this, we generated a *Slc32a1^Cre^:Ai9* mouse in which all GABAergic inhibitory neurons expressed the tdTtomato marker (Vong et al., 2011). Consistent with the Tbr2-GFP and RBPMS co-staining pattern, we found that ~54% of the Tbr2^+^ cells in the GCL are Ai9^+^ ACs (53.6 ± 1.5%, white arrowheads in Fig. 1G, 1H), and that the remaining ~46% Tbr2^+^ cells are RBPMS^+^ RGCs (46.4 ± 1.5%, yellow arrowheads in Fig. 1G, 1H). Using RBPMS and *Slc32a1^Cre^:Ai9* as pan-RGC and dAC readouts, respectively, we conclude that Tbr2^+^ RGCs account for 11.6 ± 1.0% of all RGCs, and that Tbr2^+^ dACs account for 18.1 ± 1.3% of all dACs (Fig. 1I). Moreover, we found by co-immunostaining using anti-melanopsin and RBPMS that ~6.3% (6.26 ± 1.23%) of RBPMS^+^ RGCs express melanopsin (Fig. 1J), suggesting that ~50% of Tbr2^+^ RGCs express Opn4 at detectable levels under our experimental conditions.

### Characterization of Tbr2-expressing RGCs

To determine the morphologies of Tbr2-expressing RGCs, we bred *Tbr2^CreERT2^* with the *Rosa^iAP^* mouse line to produce *Tbr2^CreERT2/+^: Rosa^iAP/+^* mice, injected tamoxifen through the intraperitoneal (IP) route, and then performed genetic sparse AP labeling on flat-mounted retinas (Fig. 2A) (Badea and Nathans, 2004, Badea et al., 2009b, Jamal et al., 2020). In agreement with the above finding that Tbr2 is expressed in 18% of dACs, we found that AP-marked retinal neurons were comprised of poly-axonal wide-field dACs (dendritic field diameter > 500 μm) amongst diverse types of RGCs with dendrites entangled within a dense and narrow band in IPL (Fig. 2B). By breeding *Tbr2^CreERT2^* into the Ai9 reporter background to dye-fill these dACs in tamoxifen-treated *Tbr2^CreERT2^:Ai9* retinas (Fig. 2C), we found that these dACs have long dendritic processes that ramify mainly in the ON laminae of the IPL (Fig. 2D, D’). This stratification pattern is consistent with GFP signal on the cross-sections of *Tbr2^TauGFP/+^* retinas (Fig. 1D-F). Whole cell current clamp recording of ~100 of these dACs found no apparent intrinsic photosensitivity, but several discernable IMP patterns (not shown), suggesting that they are not a homogenous group. By direct injection of 4-hydroxytamoxifen (4-OHT) into the vitreous space (1 μl, 0.1 mg/ml), we found that this induction regiment resulted in more labeled RGCs and less interference from the dAC-derived dendritic AP signals. In subsequent studies, we focused on characterizing Tbr2^+^ RGCs.

**Figure 2.**
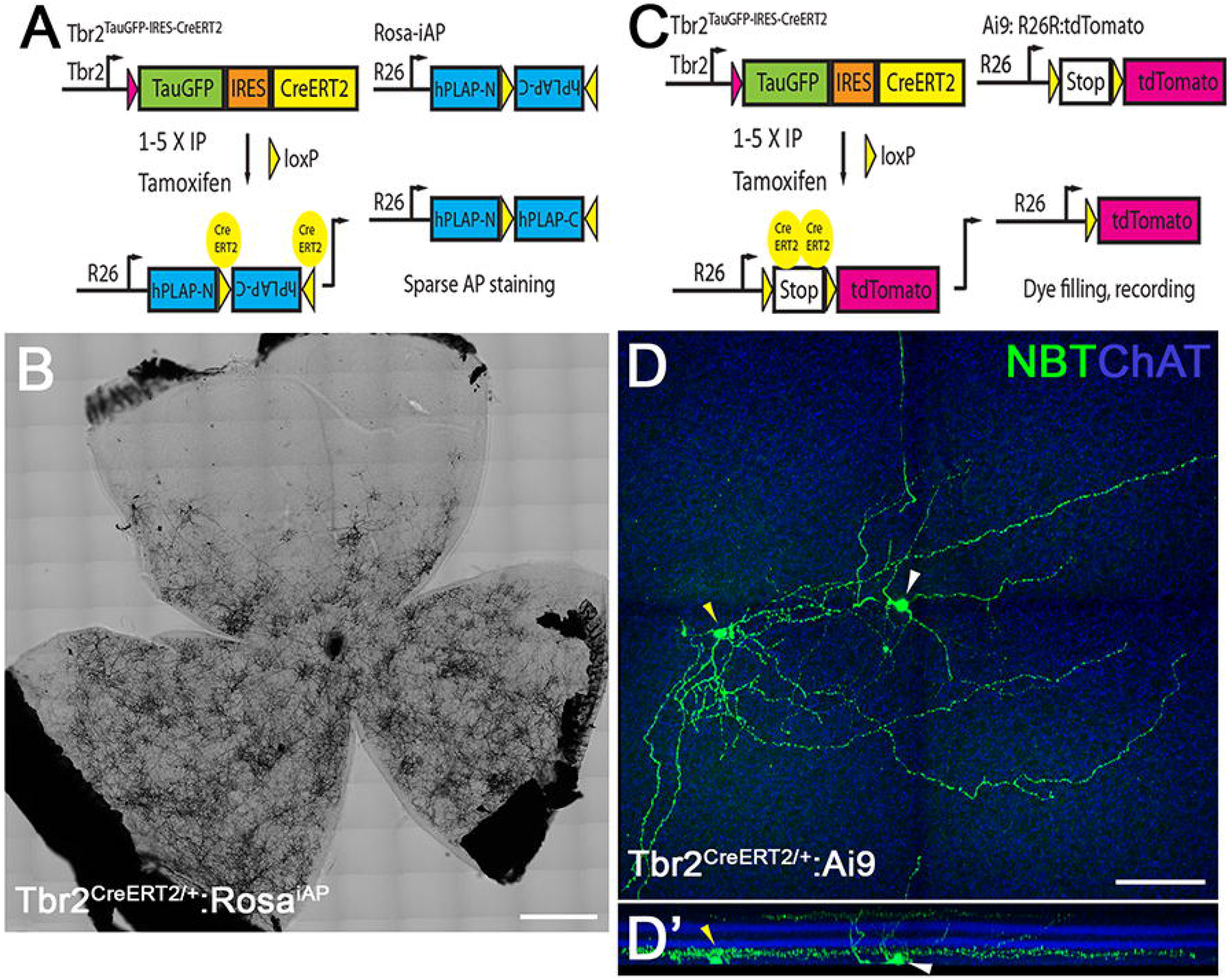
Identification of Tbr2^+^ RGCs and dACs. **(A)** The genetic sparse labeling system in the *Tbr2^CreERT2^:Rosa^iAP^* mouse line. **(B)** Representative AP staining from a flat-mounted P30 *Tbr2^CreERT2^:Rosa^iAP^* retina showing intense AP^+^ Tbr2-expressing retinal neurons. **(C)** The fluorescent labeling system of Tbr2^+^ neurons in the *Tbr2^CreERT2^:Ai9* mouse line. **(D, D’)** Representative images showing the morphology of a Tbr2-expressing displaced amacrine cell (yellow arrowhead) and a nearby RGC (white arrowhead) revealed by filled neurobiotin (D) and the location where their terminal dendrites stratify in reference to ChAT bands in IPL (D’). NBT: neurobiotin. ChAT: cholinergic acetyltransferase. Scale bars: 500 μm (B), 100 μm (D).

#### M1-like Tbr2^+^ RGCs

Among the 259 cells examined by AP staining from 29 retinas, we distinguished 44 “M1-like” cells based on their terminal dendrites stratifying to the boundary between IPL and INL, a decisive morphological feature of conventional M1 ipRGCs (Fig. 3A-3D). These cells had medium-sized somata (17.11 ±3.1 μm in diameter) but displayed large variations in dendritic arbor sizes (see green polygons in Fig. M1 A-D; arbor diameter: 269.21 ± 56.02 μm) and degree of dendritic branching (21.89 ± 6.39). The variations reflect discordant observations of M1-ipRGC sizes in the literature (Berson et al., 2010, Ecker et al., 2010, Muller et al., 2010, Schmidt and Kofuji, 2011). Most of these M1-like Tbr2^+^ RGCs have 2 or 3 primary dendrites, although we also encountered one cell with a single primary dendrite (red arrowheads in Fig. 3E, 3E’) and a smaller soma (11.3 μm vs. 17.1 μm). We also detected several Tbr2^+^ RGCs whose somata were in the INL (red arrowheads in Fig. 3F-3H). These cells have variable dendritic arbor sizes but all stratify exclusively to the OFF sub-laminae of the IPL. In agreement with our previous finding that all displaced ipRGCs express Tbr2 (Mao et al., 2014), these cells appeared to be displaced M1-ipRGCs. Because the morphological diversity suggests that these “M1-like” Tbr2^+^ RGCs may be more heterogeneous than conventional M1 ipRGCs, we then performed targeted patch clamp recording of 58 additional Tbr2^+^ cells under synaptic blockade in 8 tamoxifen-treated *Tbr2^CreERT2/+^:Ai9* retinas. We found 13 of these M1-like cells, all with intrinsic photosensitivity but inhomogeneous IMP profiles, allowing them to be divided further into three subgroups: M1n, M1r, and M1s (Fig. 3I-K). We then surveyed additional 17 such M1-like Tbr2^+^ RGCs for relative abundance and performed quantitative morphometric analysis of their dendrites. We found out of a total of 30 analyzed M1-like cells that they are about equal in abundance (36% for M1n and M1r, 28% for M1s). One-way ANOVA found no differences in primary dendrite numbers, dendritic lengths, dendritic field sizes and branching numbers among them but M1s has a much lower input resistance at 463.7 ± 90.5 MΩ than those of M1n and M1r at 1259.9 ± 110.1 MΩ and 1252.8 ± 109.7 MΩ, respectively. Given a recent finding that M1-ipRGCs possess a wide range of photosensitivity (Milner and Do, 2017), the exact relationship between input resistance and intrinsic photosensitivity among these three M1-like Tbr2^+^ cells will be important to determine in future studies.

**Figure 3.**
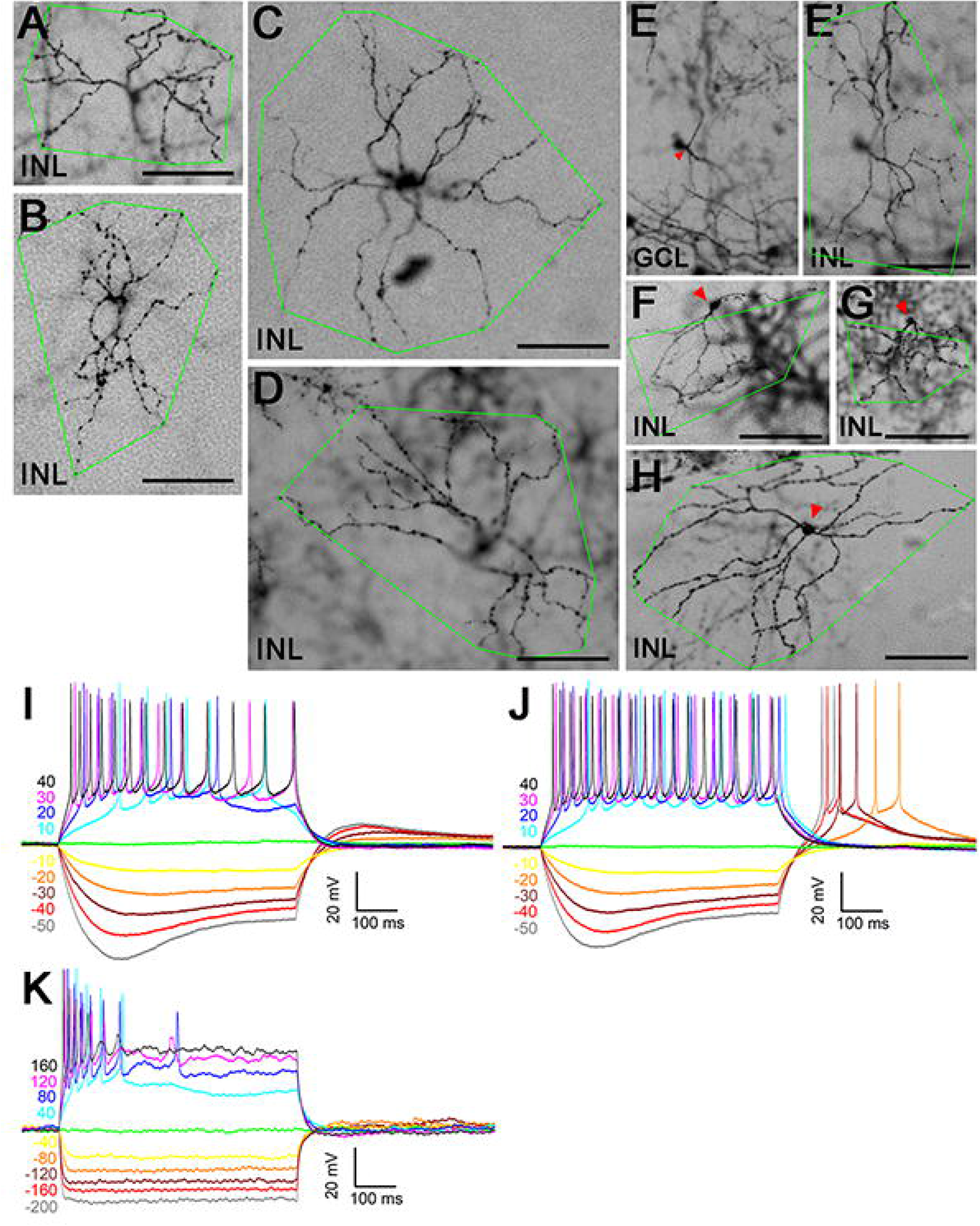
M1-like Tbr2^+^ RGCs. **(A-H)** Representative brightfield images of AP-stained *Tbr2^CreERT2/+^:Rosa^iAP/+^* retinal flatmounts. Images were focused on the terminal dendrites located in between INL/IPL boundary (labeled as INL) except in (E) where the focus was on the soma in the ganglion cell layer. The green polygons represent the terminal dendritic arbor. The red arrowhead in E indicates the single primary dendrite of this RGC. Three representative displaced M1-like RGCs (F-H) with somata located in INL (pointed by red arrowheads) were also shown. (I-K) Representative intrinsic membrane property (IMP) profiles from three M1-like Tbr2^+^ RGC subtypes: M1n, M1r, and M1s. Membrane potential changes in response to 600 msec current injecteions of varying amplitudes and polarities as indicated at left. Note the larger currents required to elicit changes in M1s cells. GCL: ganglion cell layer. INL: inner nuclear layer. IPL: inner plexiform layer. Scale bars: 100 μm (A-H).

#### Bistratified Tbr2^+^ RGCs

We identified 58 AP-labeled bistratified RGCs with diverse morphological characteristics (Fig. 4). The first type (n=12) had smaller dendritic arbor sizes and extensive branching patterns (Fig. 4A, 4B). Their 2 dendritic arbors stratified next to the inner (OFF) and outer (ON) margins in the IPL, and the dendritic arbors in the ON sub-laminae (red polygons) were larger than that in the OFF sub-laminae (green polygons) (Fig. 4A-4B’). This type of RGC is similar to the recently described M6 ipRGCs from a Cdh3-GFP mouse line (Quattrochi et al., 2019). The other 46 bistratified Tbr2^+^ RGCs had relatively larger and less branching dendritic arbors (Fig. 4C-D’). Nineteen of these cells had dendrites confined almost entirely to the ON sub-laminae (Fig. 4C, 4D) with only a few straying ones ramifying up into the OFF sub-laminae (black arrowheads in Fig. 4C’, 4D’). Thirteen cells had similar dendritic arbors in ON and OFF sub-lamina (Fig. 4E, 4E’), while the remaining 14 cells had dendrites stratified mainly to the OFF-layer (Fig. 4 F, 4F’). These larger bistratified RGCs are similar to the M3 ipRGCs described previously (Schmidt and Kofuji, 2011). We initially encountered 4 intrinsically photosensitive bistratified Tbr2^+^ RGCs by whole cell recording under synaptic blockade, including one M6-like cell that was responsive only under the brightest stimulation at 5.9 log photon/μm^2^. Subsequent survey of 8 additional bistratified Tbr2^+^ RGCs showed that they have diverse IMP profiles (Fig. 4G-J). These bistratified Tbr2^+^ RGCs therefore appear to also be diverse in dendritic morphology, IMP, and likely in their intrinsic photoresponses.

**Figure 4.**
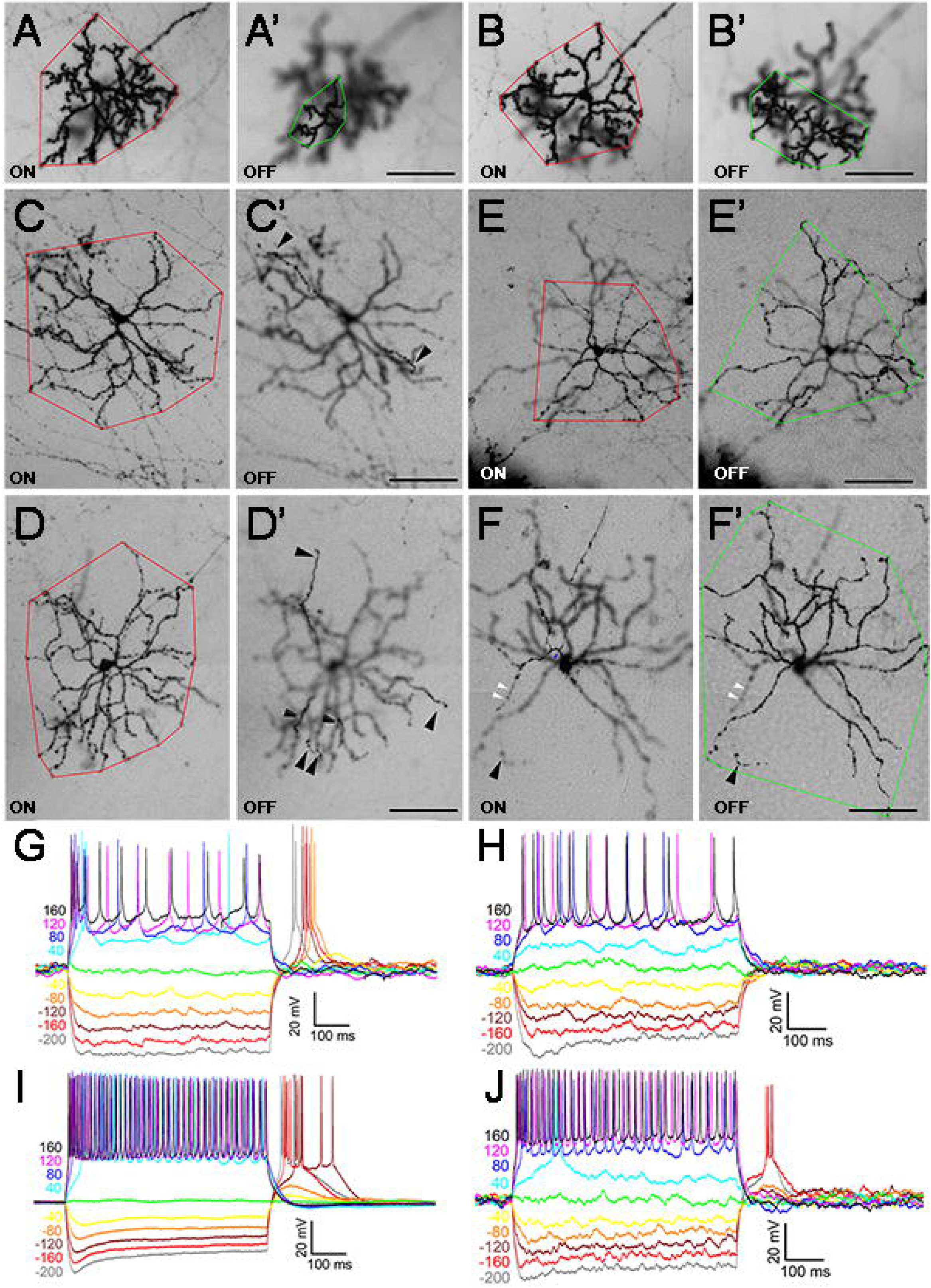
Bistratified Tbr2^+^ RGCs. **(A-F’)** Representative AP-stained images of M6-like **(A-B’)** and various types of M3-like **(C-F’)** Tbr2^+^ RGCs from *Tbr2^CreERT2/+^:Rosa^iAP/+^* retinal flatmounts. Images were focused on the ON (A-F) or OFF (A’-F’) IPL sub-lamina.. Red polygons represent the terminal dendritic arbor in the ON-layer, while the green polygons represent those in the OFF-layer. The white arrowheads indicate the sparse terminal dendrites in the ON layer, and the black arrowheads indicat those in the OFF-layer. (G-J) Representative and diverse IMP profiles associated with the bistratified Tbr2^+^ RGCs. The M6 subtype has a stereotypic IMP profile (J), the M3 subtypes have not. Scale bars: 100 μm (A’-F’).

#### ON-layer stratified Tbr2^+^ RGCs

Because their dendrites were often surrounded by the massive dendritic network from other RGCs and dACs, detailed analysis were done on 157 AP-stained RGCs not entrenched with others. Their dendritic stratification appeared only in the ON sub-laminae with dendritic characteristics similar to published ON-layer stratified M2, M4, and M5 ipRGC subtypes. Below we separate them by sizes of their dendritic arbors, somata, branching points and IMPs.

#### M4-like Tbr2^+^ RGCs

M4 ipRGCs have been characterized as having the largest dendritic arbor areas and soma diameters among all known ipRGCs (Estevez et al., 2012) and a recent study identified them intersectionally in *Opn4^Cre^:Z/EG* mice by coincident GFP and SMI32 signals (Sonoda et al., 2019). In 29 AP-stained retinas, we analyzed 13 these M4-like Tbr2^+^ RGCs with large dendritic arbors and somata (arbor diameter: 320.48 ± 50.5 μm; soma diameter: 19.54 ± 3.72 μm; N=13) and relatively loose dendritic arbors (Fig. 5A-5B’). They displayed less complex branching (branch points: 28.67 ± 2.53) when compared with other ON-layer stratified Tbr2^+^ RGCs. We encountered 3 of these M4-like cells by whole cell recording and all of them display a characteristic IMP profile previously described (Kiyama et al, 2019; Fig. 5C). Interestingly, one of them exhibited no intrinsic photosensitivity under our brightest stimulation.

**Figure 5.**
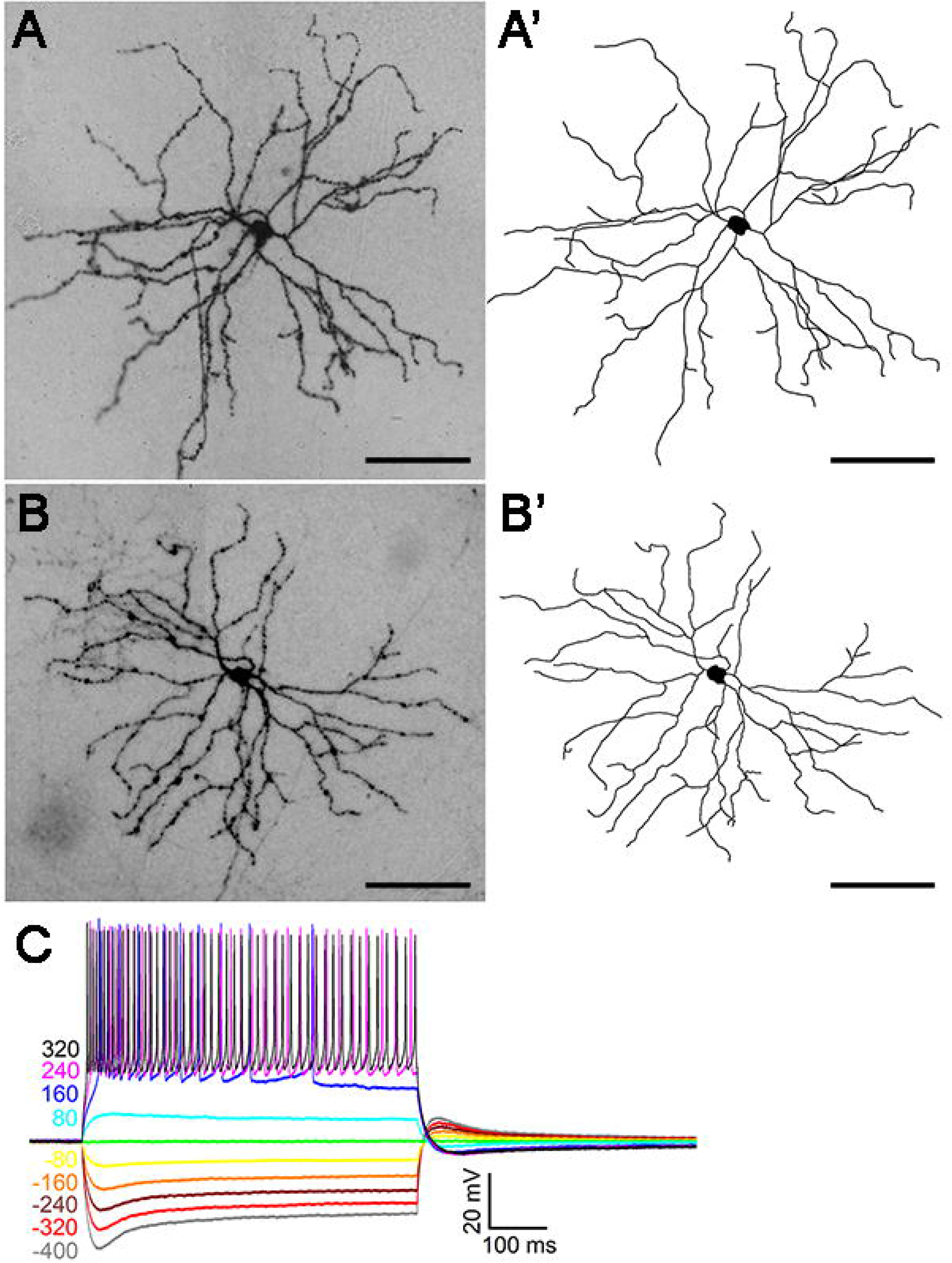
Large M4-like Tbr2^+^ RGCs. **(A-B)** Representative images of AP-stained M4-like Tbr2^+^ RGCs from *Tbr2^CreERT2/+^:Rosa^iAP/+^* retinal flatmounts. **(A’-B’)** Neuronal tracing of the ministacked images shown in A-B. (**C**). Stereotypic IMP profile of M4-like Tbr2^+^ RGCs. Scale bars: 100 μm (A-B’).

#### M5-like Tbr2^+^ RGCs

The M5 ipRGC subtype has dense, bushy dendritic trees with smaller dendritic arbors compared to M2 and M4 types and stratifies below the ON ChAT band in the IPL (Ecker et al., 2010, Stabio et al., 2018). We found 30 AP-stained Tbr2^+^ RGCs with these M5 morphological characteristics (arbor diameter: 157.28 ± 25.02 μm; soma diameter: 14.48 ± 1.93 μm, N=30, Fig. 6A, B). Notably, their dendrites emanated from 3 to 5 primary dendrites (3.77 ± 0.72, N=22) and stratified slightly below the ON ChAT band (depth in IPL: 85.73% ± 1.48%, N=20), forming more extensive branching than other ipRGCs (49.33 ± 9.68, N=15). Similar to the AP-based survey, we frequently encountered these M5-like Tbr2^+^ cells during whole cell recording with some displayed weak intrinsic photosensitivity, while others unresponsive to our brightest stimulation (not shown). The IMP profiles of these M5-like Tbr2^+^ cells appeared diverse but all contained a unique characteristic delayed return to baseline following positive current injections (Fig. 6C-E, arrows).

**Figure 6.**
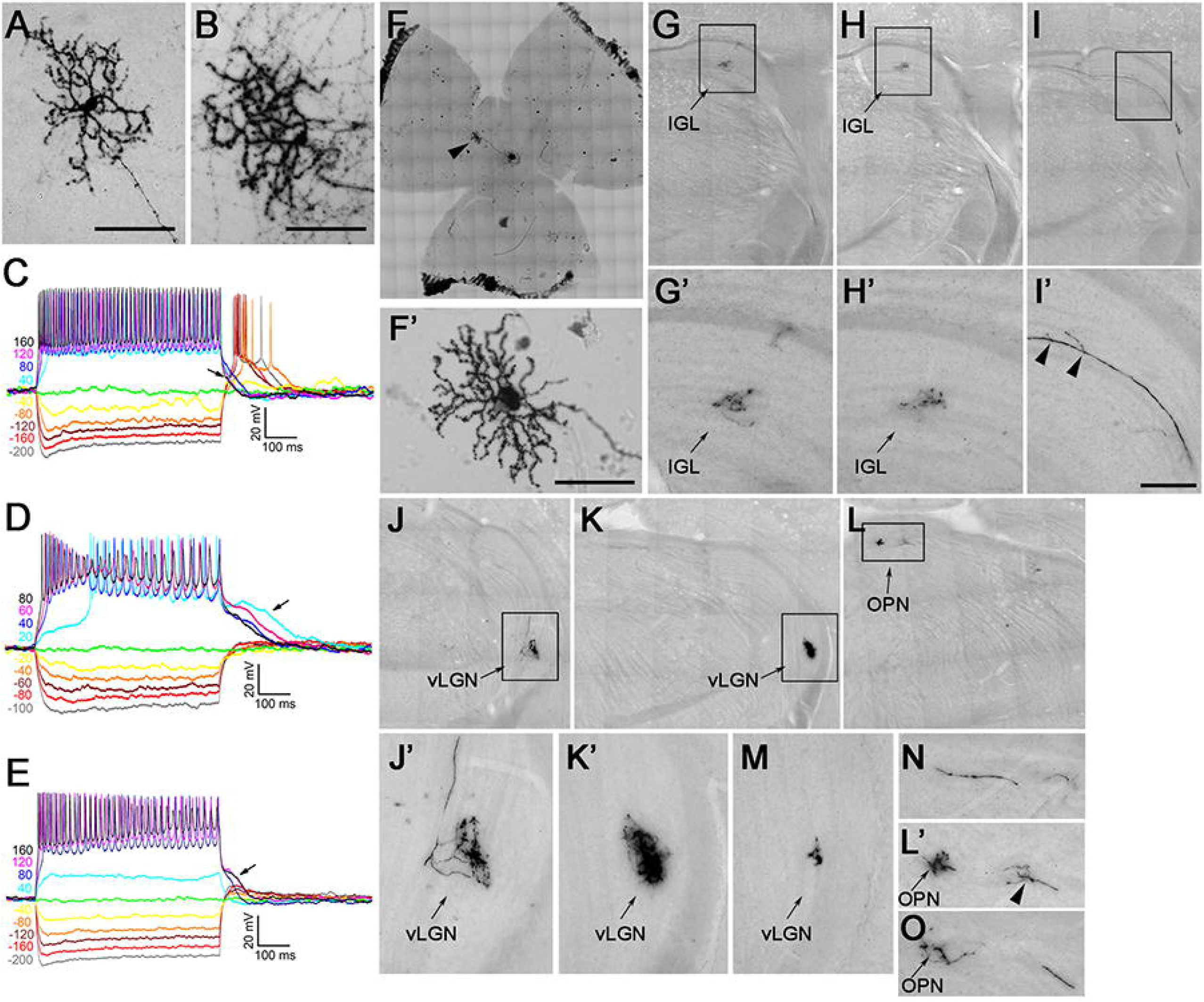
Small M5-like Tbr2+ RGCs. **(A-B)** Representative AP-stained images of M5-like Tbr2^+^ RGCs from *Tbr2^CreERT2/+^:Rosa^iAP/+^* retinal flatmounts. (**C-E**) Representative and diverse IMP profiles of M5-like Tbr2^+^ RGCs. Note the characteristic return delay to the baseline following positive current injections (arrows). **(F-F’)** Retina with a single M5-like Tbr2^+^ RGC. **(G-O)** AP-stained brain sections showing multiple axonal projections of the M5-like RGC shown in **F**’ in several different regions in the brain. **(G’-L’)** Enlarged images of the bracketed regions in G to L, respectively. IGL: inter-geniculate leaflet. OPN: olivary pretectal nuclei. vLGN: ventral lateral geniculate nuclei. INL: inner nuclear layer. Scale bars: 100 μm (A, B, F’).

Fortuitously, in one animal whose right eye was injected with a low dose of 4-OHT (1 μg/μl), we found a single M5-like RGC in the entire retina (Fig. 6F, 6F’). AP staining on brain sections revealed its retinofugal projection to the IGL (Fig. 6G-6H’), vLGN (Fig. 6J-6K’,6M), and OPN (Fig. 6L, 6L’, 6N). Notably, the size and complexity of its axonal arbors were different in different brain regions. In addition, the axon bifurcated and gave rise to several telodendria along its path (arrowheads in Fig. 6I’, 6L’). Such projection patterns are similar to those from a single M1-ipRGC, which sends collateral axonal inputs to SCN, IGL, and other brain regions (Fernandez et al., 2016). Whether this is a common feature of all ipRGCs will be an interesting topic for future investigation.

#### M2-like Tbr2^+^ RGCs

M2 ipRGCs have an intermediate dendritic arbor size between M4 and M5. We analyzed 14 M2-like AP-stained Tbr2^+^ RGCs (Fig. 7A-7D’). They had relative medium dendritic arbors (arbor diameter: 228.05 ± 28.77 μm; N=13) and somata (soma diameter: 16.44 ± 2.48 μm; N=12). Their dendrites emanated from 3 to 5 primary dendrites (3.87 ± 0.69, N=15) and branched more variably than other ipRGCs (37.08 ± 11.24, N=14). These M2-like cells possessed an IMP profile different from those of M4-like and M5-like cells, characterized by with the lack of rebound depolarization and delayed return to baseline following current injections (Fig. 7D). We encountered only two such M2-like Tbr2^+^ cells and both were intrinsically photosensitive.

**Figure 7.**
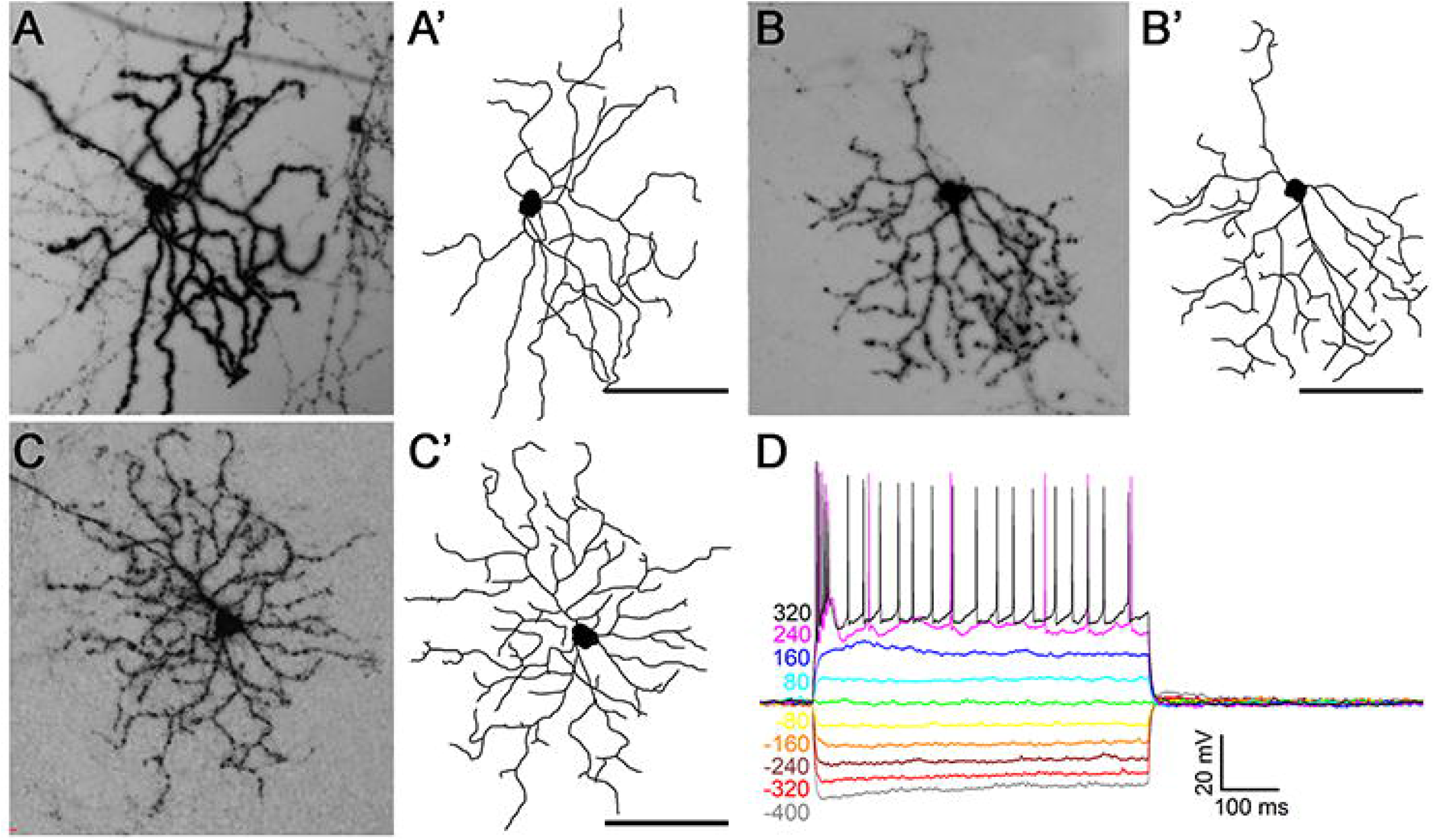
M2-like Tbr2^+^ RGCs. **(A-C)** Representative AP-stained images of 3 M2-like Tbr2^+^ RGCs from *Tbr2^CreERT2/+^:Rosa^iAP/+^* retinal flatmounts. **(A’-C’)** Corresponding tracing of ministacked images shown in A to C. (**D**) Representative stereotypic IMP profile of M2-like Tbr2^+^ RGCs. Note the quick return to baseline and the absence of regenerative changes following large current injections. Scale bars: 100 μm (A’-C’).

#### Pou4f1-expressing Tbr2^+^ RGCs

A small percentage of Tbr2-expressing cells expressed RGC marker Pou4f1 (Fig. 8A), which is not expressed, or at a very low level, in ipRGCs (Sweeney et al., 2014), suggesting that these cells are different from aforementioned reservoir ipRGCs. To track them, we bred *Tbr2^CreERT2^* and *Pou4f1^CKO^* to produce a *Tbr2^CreERT2/+^:Pou4f1^CKO/+^* mouse line for sparse AP staining (Fig. 8B). In each retina we examined, we detected 0 to 2 AP-positive RGCs. The most notable morphological characteristics was their dense and bushy dendrites (Fig. 8C-8H’) resulting from numerous branching with short terminal dendrites emanating from several primary dendrites (4.15 ± 0.36, N=13). Their dendritic arbors and soma sizes were relatively smaller (dendritic arbor diameter: 196.5 ± 26.8 μm; soma diameter: 15.54 ± 1.1 μm; N=14) and their terminal dendrites stratified to the OFF sub-laminae (depth in the IPL: 12.8% ± 1.5%, N=12). We classified this RGC type into a unique but rare Tbr2^+^Pou4f1^+^ RGC subtype because we could only detected 3 of them in all *Tbr2^CreERT2/+:^Rosa^iAP^* retinas examined. We did not encounter this rare type during whole cell recording.

**Figure 8.**
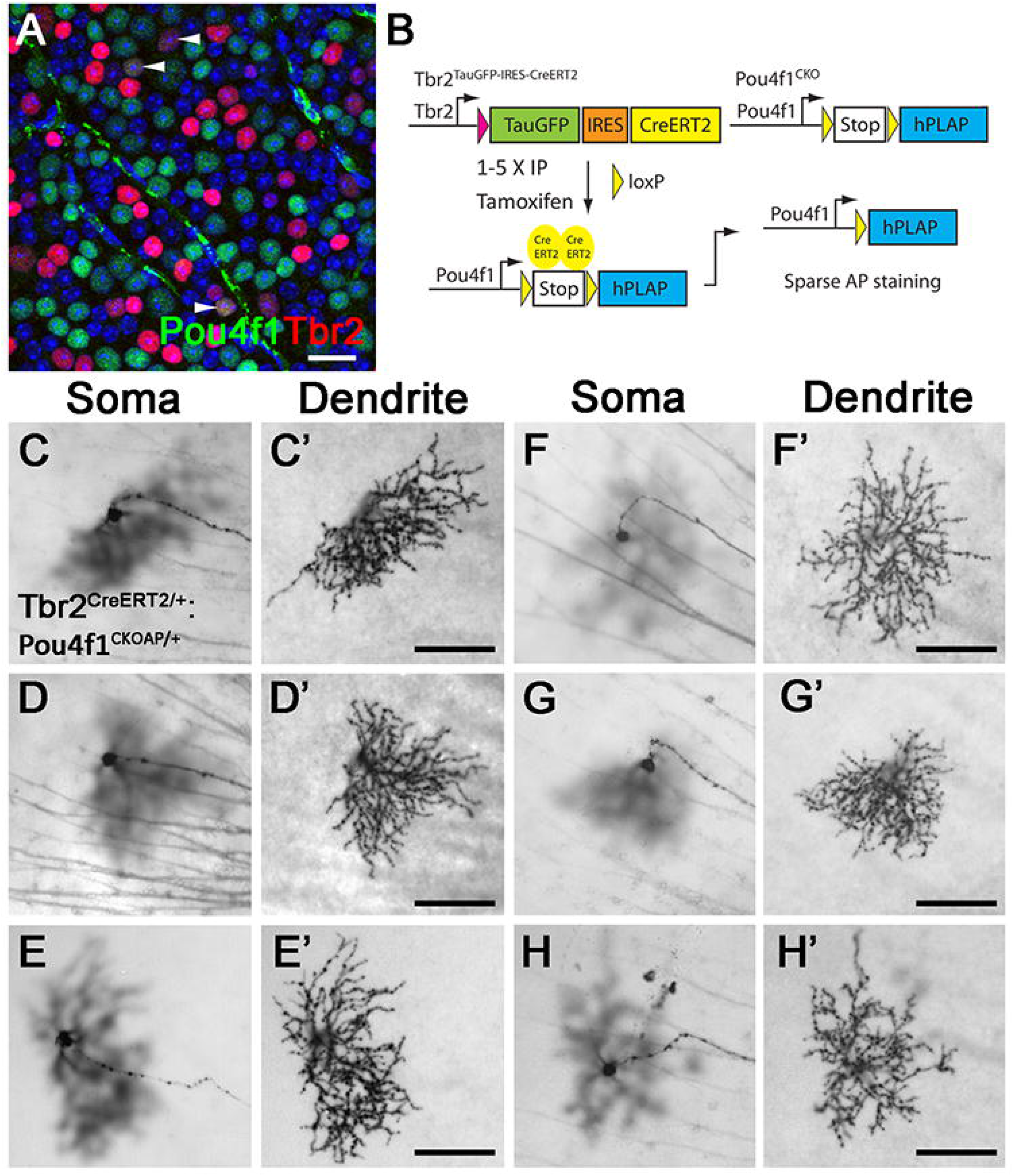
OFF-layer stratified Tbr2^+^Pou4f1^+^ RGCs. **(A)** Fluorescent image from a representative wild-type retinal flatmount showing Tbr2 expression (red) with Pou4f1-expressing (green) RGCs (white arrowheads). **(B)** The genetic sparse labeling system in the *Tbr2^CreERT2^:Pou4f1^CKO^* mouse line for brightfield identification of this cell type by AP staining. **(C-H’)** Representative brightfield AP-stained images of the Tbr2^+^Pou4f1^+^ RGCs. Scale bars: 100 μm (C’-H’). AP: alkaline phosphatase.

### Axonal projection of Tbr2^+^ RGCs in brain

To determine the central projection of Tbr2^+^ RGCs, we examined AP staining patterns on brain sections from *Tbr2^CreERT2/+:^Rosa^iAP/+^* mice that received 5 consecutive tamoxifen doses through the IP route. As expected, intensive AP staining was detected in the SCN (Fig. 9A), vLGN, IGL, and OPN (Fig. 9B-9C). In addition, A few AP^+^ axonal arbors with relatively diffused signals were found in the dorsal lateral geniculate nuclei (dLGN) (Fig. 9B). These data confirm that Tbr2^+^ RGCs are not merely morphologically similar to ipRGCs, but they also have identical retinofugal projections to known ipRGCs.

**Fig. 9.**
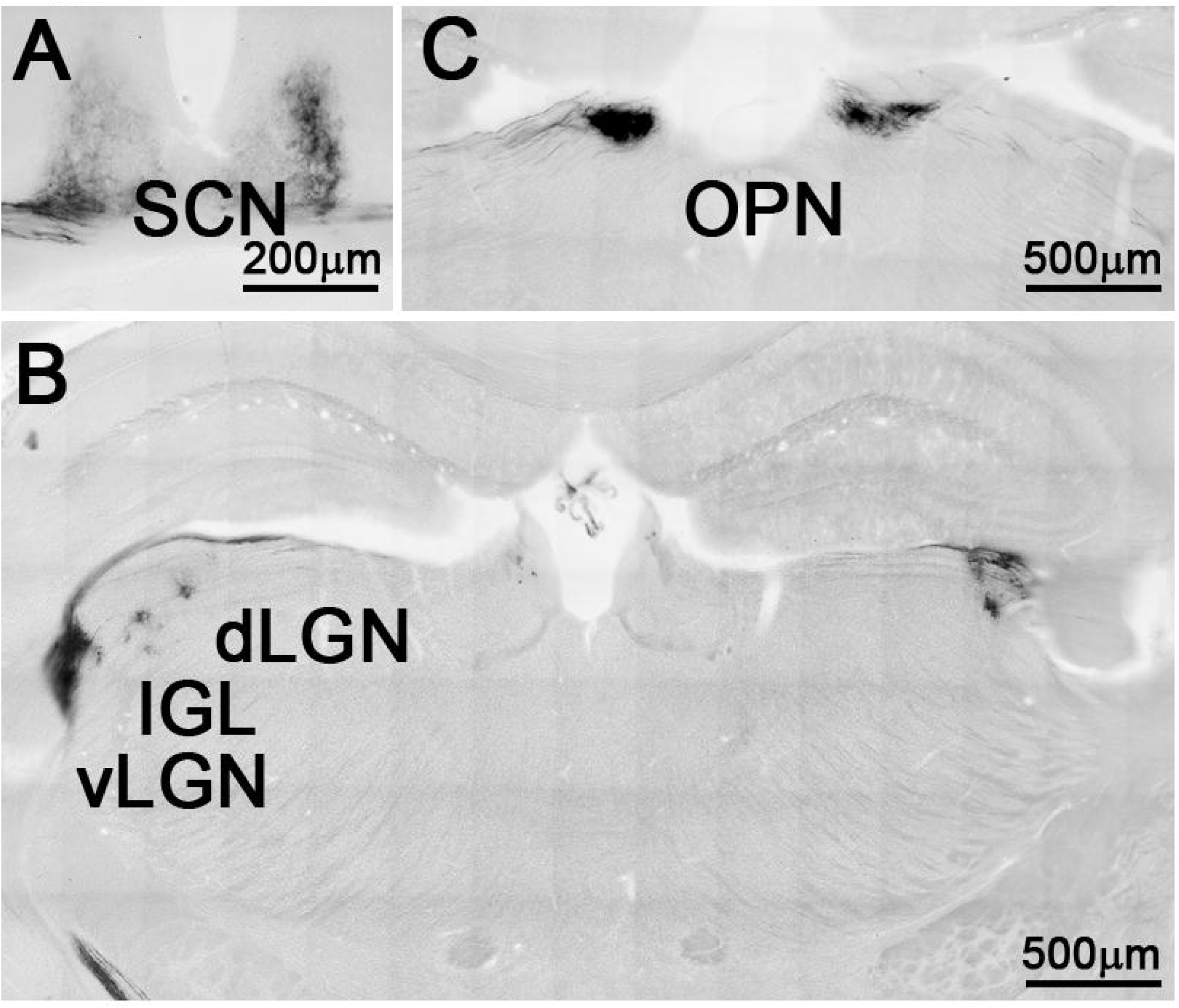
Retinofugal projections of Tbr2^+^ RGCs in *Tbr2^CreERT2/+^:Rosa^iAP^* mice. **(A-C)** AP-stained brightfield images showing projections of Tbr2^+^ RGCs into SCN (A), IGL, vLGN, dLGN (B), and OPN (C). IGL: inter-geniculate leaflet. vLGN: ventral lateral geniculate nuclei. dLGN: dorsal lateral geniculate nuclei. OPN: olivary pretectal nuclei. SCN: suprachiasmatic nuclei.

Next, we attempted to resolve a standing discrepancy between our previous study and another study on whether all SCN-afferent RGCs express melanopsin (Baver et al., 2008). To do this, we compared the projection patterns of Tbr2^+^ RGCs in the control *Tbr2^TauGFP/+^* mouse (Fig. 10A) and *Tbr2^TauGFP/fx^:Opn4^Cre/+^* mouse line in which native ipRGCs are eliminated but Tbr2^+^ ipRGC reservoir neurons are not (Fig. 10F) (Mao et al., 2014). We injected Alexa 555-cholera toxin subunit B (CTB-555) into the right eyes of these mice to trace RGC axonal targets in brains and then compared the GFP and CTB-555 signals on brain sections. We found that GFP^+^ cells from mice with reservoir-only Tbr2^+^ cells projected to the same regions as those from control mice in the SCN (Fig. 10C’, 10H’), vLGN and IGL (Fig. 10D’, 10I’), and OPN (Fig. 10E’, 10J’) and that GFP signals in the control were much higher than those in the mutant (compare Fig. 10C’ and 10H’, 10D’ and 10I’, and 10E’ and 10J’) where most of the native ipRGCs have been eliminated. The weak GFP signal can also be seen in the dLGN (Fig. 10C’, 10H’) but no GFP signal was found in the superior colliculus and the accessory optic system (not shown). Together, these data confirmed that Tbr2^+^ RGCs projected to the same region as ipRGCs. The number of GFP^+^ cells in the *Tbr2^TauGFP/flox^:Opn4^Cre/+^* retinas was reduced significantly compared to the *Tbr2^TauGFP/+^* control retinas (compare Fig. 10B and 10G), again supporting the notion that Tbr2^+^ RGCs are ipRGCs and express Opn4 sometime in their lifetime.

**Figure 10.**
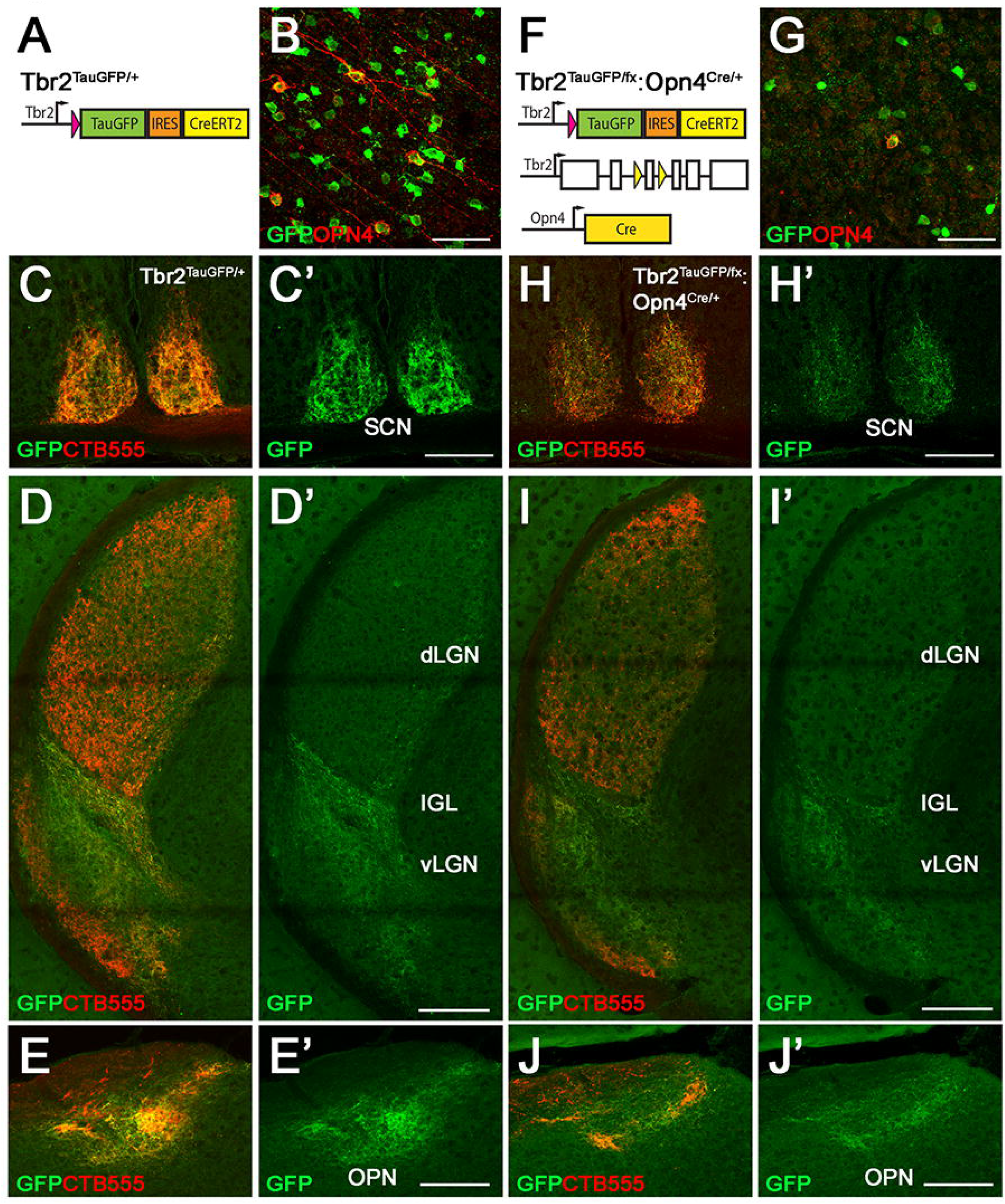
Identical retinofugal projections between Tbr2^+^ RGCs and reservoir Tbr2^+^ ipRGCs in the brain. **(A-E’)** Fluorescent images from GFP-stained retinas and brain sections of *Tbr2^TauGFP/+^* mice with right eyes injected with CTB-555 (A). Representative fluorescent image showing GFP signals (green) and melanopsin (red) in the retina (B). Intense GFP signals evident in SCN (C, C’), vLGN (D, D’), and OPN (E, E’), and only sparse and relatively weak GFP signals in dLGN. **(F-J’)** Fluorescent images from GFP-stained retinas and brain sections of *Tbr2^TauGFP/fx^:Opn4^Cre/+^* mice with right eyes injected with CTB-555 (F). Co-immunostaining showing GFP signals (green) and melanopsin (red) in retina (G). Clearly noticeable but weaker GFP signals were detected in SCN (H, H’), vLGN (I, I’), and OPN (J, J’). OPN: olivary pretectal nuclei. SCN: suprachiasmatic nuclei. dLGN: dorsal lateral geniculate nuclei. vLGN: ventral lateral geniculate nuclei. IGL: inter-geniculate leaflet. Scale bars: 50 μm (B, G), 200 μm (C’–E’, H’–J’).

### Eliminating Tbr2-expressing RGCs by melanopsin-SAP

We previously speculated that Tbr2^+^ RGCs include ipRGC reservoir neurons with very low OPN4 expression level or they do not express Opn4 until the death of native ipRGCs (Mao et al., 2014). We reasoned that, in either case, melanopsin-SAP (an anti-melanopsin antibody conjugated with a ribosome-inactivating protein saporin), which could effectively kill melanopsin^+^ ipRGCs (Goz et al., 2008), should eliminate most if not all Tbr2^+^ RGCs. We injected melanopsin-SAP (1 μl, 400 ng/μl) into one eye of *Tbr2^TauGFP/+^* mouse and rabbit IgG-SAP into the other as a negative control (1 μl, 400 ng/μl), then isolated the retinas 30 days later to check for GFP and melanopsin signals. We found in our first trial that approximately 33% of Tbr2^+^ cells in the ventral region were abolished, but this effect was less prominent in the other 3 quadrants (data not shown). We then tested multiple injections of melanopsin-SAP (one injection per month, total 2 injections) and isolated retinas 30 days after the second injection and found that approximately 41% of Tbr2^+^ cells were abolished (Ctrl: 643.7 ± 106.7; melanopsin-SAP: 380.3 ± 47.1, P = 0.0174, per image area 1.225 x 10^5^ μm^2^) with few remaining melanopsin^+^ cells (Fig. 11A-11D’). These data indicate that the majority of Tbr2^+^ RGCs indeed express melanopsin at a level not detectable by conventional immunostaining but sufficient to be eliminated by melanopsin-SAP.

**Figure 11.**
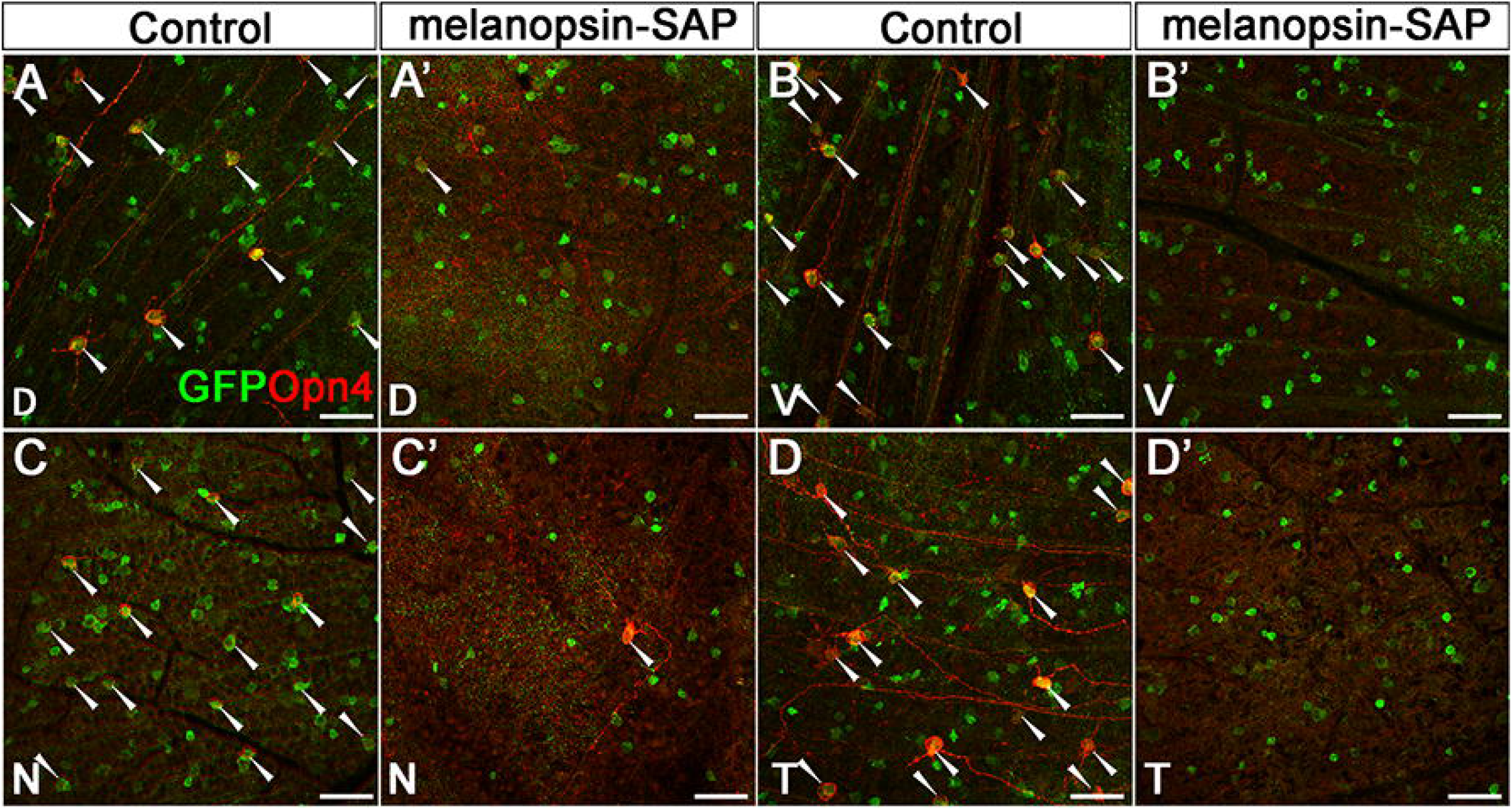
Ablation of Tbr2-expressing RGCs by melanopsin-SAP. (**A-D’**) Co-immunofluorescent images stained by anti-GFP and anti-melanopsin antibodies on *Tbr2^TauGFP/+^* retinas that were treated with rabbit-IgG-SAP (A–D) or anti-melanopsin-SAP (A’–D’). D: dorsal, V: ventral, N: Nasal, T: temporal areas of retinas. Scale bar: 50 μm (A-D’).

### *Tbr2* is essential for the survival of Tbr2^+^ RGCs and *Opn4* expression in adult retina

Previously we examined *Opn4^Cre^:Tbr2^fx/fx^* retinas and found that most ipRGCs are abolished, leading to the conclusion that *Tbr2* is essential for maintaining the survival of ipRGCs. However, *Opn4* is expressed as early as E14, leaving a caveat that the phenotype might result from the requirement of *Tbr2* for the survival of ipRGCs during development. To test whether Tbr2 is essential for ipRGC survival in adult retinas, we injected AAV2-Cre into the vitreous space of adult (P30) *Tbr2^TauGFP/+^* (control) and *Tbr2^TauGFP/fx^* mice, isolated retinas 12 days post-injection, and examined co-immunostaining patterns of Tbr2, GFP, and RBPMS. We found that while *Tbr2* expression was efficiently removed by AAV2-Cre in *Tbr2^TauGFP/fx^* retinas after 12 days, many GFP^+^ Tbr2-deficient RGCs remained (double white arrowheads in Fig. 12B). However, at 21 days post-injection, we found that GFP^+^RBPMS^+^ cells in AAV2-Cre treated *Tbr2^TauGFP/fx^* retinas decreased by approximately 50% (data not shown). At 38 days post-injection, we could barely find GFP^+^RBPMS^+^ RGCs in *Tbr2*-deleted regions (compared Fig. 12C and 12D. Ctrl: 55.2 ± 4.8; AAV2-Cre: 10.2 ±_1.5, P = 2.2E-06, per unit area 1.0 x 10^5^ μm^2^). Together, these data indicate that *Tbr2* is essential for the survival of Tbr2^+^ reservoir ipRGCs and dACs in adult retinas, supporting our previous conclusion.

**Figure 12.**
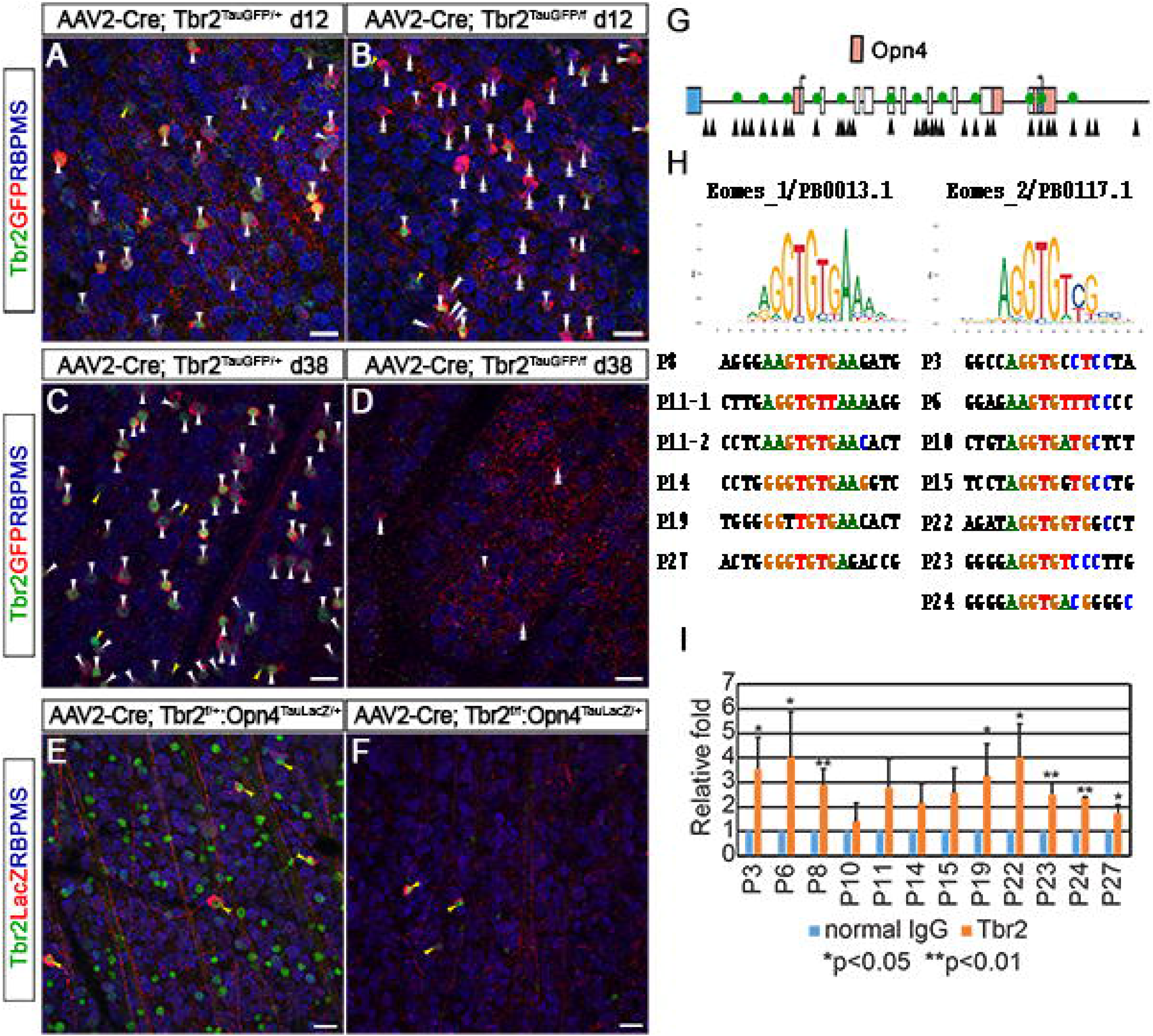
Tbr2 is essential for maintaining ipRGC survival and Opn4 expression. **(A-D)** Fluorescent images of Tbr2, GFP, and RBPMS on *Tbr2^TauGFP/+^* retinas (A, C) and *Tbr2^TauGFP/fx^* (B, D) that was treated with AAV2-Cre. Retinas were isolated 12 days (A, B) or 38 days (C, D) post-injection. **(E-F)** Co-immunofluorescent staining of Tbr2, LacZ, and RBPMS on *Tbr2^fx/+^:Opn4^TaulacZ^* (E) or *Tbr2^fx/fx^:Opn4^TaulacZ^* (F) retinas. Retinas were isolated 14 days post-injection. **(G)** Schematic illustration showing the presumptive Tbr2-binding elements (arrowheads) in *Opn4* locus (brown boxes). The green circles indicate the confirmed Tbr2 binding sites by ChIP-qPCR. **(H)** Alignment of the validated Tbr2-bound sequences in *Opn4* with the two conserved Eomes binding sequences, Eomes_1 and Eomes_2, in Jaspar (http://jaspar.genereg.net/). **(I)** ChIP-qPCR analysis of WT retinas (P0) were performed to measure relative antibody-bound DNA fragments of the *Opn4* region. Each bar represents the average result from 3 independent ChIP experiments. Scale bars: 20 μm (A-F). ChIP-qPCR: Chromatin Immunoprecipitation (ChIP) coupled with quantitative PCR.

Because there was a time window between the deletion of *Tbr2* (12 days post-injection) and the disappearance of Tbr2-deficient RGCs (38 days post-injection), we reasoned that if *Opn4* expression is regulated by Tbr2, *Opn4* should be down-regulated while the Tbr2-deficient RGCs were still alive between P12 and P38. To test this, we injected AAV2-Cre into the vitreous space of P21 *Tbr2^fx/+^:Opn4^TauLacZ/+^* (control) and *Tbr2^fx/fx^:Opn4^TauLacZ/+^* retinas, then isolated retinas 14 days post-injection and conducted co-immunostaining for Tbr2, RBPMS, and Opn4-driven LacZ. We detected fewer LacZ-expressing ipRGCs in the *Tbr2-deleted* region in *Tbr2^fx/fx^:Opn4^TauLacZ/+^* retinas (compared Fig. 12E and 12F. Ctrl: 6.3 ± 0.6; AAV2-Cre: 3 ± 0.7, P = 0.0003, per unit area 1.0 x 10^5^ μm^2^). Additionally, most of the remaining LacZ^+^ cells expressed Tbr2, suggesting that these cells were not infected by AAV2-Cre. Together, these data suggest that *Tbr2* is essential for maintaining *Opn4* expression, likely through direct regulation.

To test whether Tbr2 directly regulates Opn4 expression, we sought to test whether it binds to *cis*-regulatory *T*-elements present in the *Opn4* locus. We conducted JASPAR *in-silico* analysis in the *Opn4* locus (9.6 kb gene body ± 4kb) and found 33 candidate Tbr2-binding *T*-elements (arrowheads in Fig. 12G) (Khan et al., 2018). Using an anti-Tbr2 antibody from Abcam (#ab23345) previously known to pull down Tbr2-bound chromatin from mouse cortical neurons (Sessa et al., 2017), we conducted chromatin immunoprecipitation (ChIP) from a pool of 20 P0 WT retinas for ChIP-qPCR, and surprisingly we found that Tbr2 occupied 13 of these presumptive *T*-elements at varying levels (green circles in Fig. 12G, 12H, 12I). These data suggest a direct regulatory role of Tbr2 on *Opn4* transcription.

## Discussion

*Tbr2* is an important transcription factor essential for many developmental processes including trophoblast development, mesoderm formation, T cell differentiation, induction of anterior visceral endoderm, and neurogenesis of olfactory and cortical neurons. (Russ et al., 2000, Pearce et al., 2003, Arnold et al., 2008, Sessa et al., 2008, Sessa et al., 2010, Mizuguchi et al., 2012, Nowotschin et al., 2013). In retina, we were the first to show that it is a direct downstream target of Pou4f2, implicating its role as a developmental regulator for RGC subtypes within the hierarchical transcription network controlling RGC development (Mao et al., 2008, Mu et al., 2008). In subsequent 2 independent studies, *Tbr2* was shown to be essential for ipRGC formation (Sweeney et al., 2014, Mao et al., 2014). Using genetic loss-of-function strategies, we further demonstrated that *Tbr2* is indispensable for ipRGC maintenance and hence proposed that *Opn4* expression may be activated in Tbr2^+^ RGCs, allowing them to serve as a reservoir for ipRGCs (Mao et al., 2014). In this study, we developed new mouse resources to address a number of unanswered questions relating to this hypothesis.

### Tbr2^+^ RGCs are ipRGCs

The first and foremost puzzle is why are there so many Tbr2^+^ cells but only a small fraction express detectable level of *Opn4?* Our current co-immunostaining and genetically sparse labeling studies reveal that approximately 54% of Tbr2-expressing cells in the GCL are not RGCs but dACs and that the Tbr2^+^ dACs account for approximately 18% of all dACs in the GCL. The identities and functions of these apparently diverse dACs were not determined here but In the RGC population, we found that Tbr2^+^ RGCs account for approximately 12% of all RGCs, much less than our previous estimation. The previous assumption that all Tbr2^+^ cells were RGCs must therefore be discarded. Moreover, using melanopsin expression together with a pan-RGC marker (RBPMS), we found that melanopsin-expressing cells account for approximately 6% of all RGCs. This number now approximates the number of GFP^+^ RGCs found in *Opn4^Cre^:Z/EG* retinas in our previous analysis (No. of GFP^+^: 2028 at 1-month-old vs. 2870 at 10 months) (Mao et al., 2014). However, it still represents only about half of all Tbr2^+^ RGCs. Based on the current findings that Tbr2^+^ RGCs are anatomically indistinguishable from known ipRGC types and with the same retinofugal projections and most of them possess varying degrees of intrinsic photosensitivity, we have strengthened the notion that Tbr2^+^ RGCs are indeed reservoir ipRGCs. The simplest reconciling explanation is that some Tbr2^+^ RGCs express *Opn4* at detectable levels by conventional immunostaining and those whose levels fall below detection were not accounted for. One solid line of evidence supporting this explanation is that the melanopsin-SAP ablated nearly 40% of Tbr2^+^ cells (Fig. 11). Another supporting evidence is that the majority of Tbr2^+^ RGCs we recorded exhibited intrinsic light sensitivity, as whole-cell recording is more sensitive than immunostaining in revealing Opn4 expression. Another more sophisticated explanation, according to the literature, is that the expression of *Opn4* within the entire Tbr2^+^ RGC population is dynamically regulated by ambient light level in the environment. It is known in rat that ipRGC number and *Opn4* level can be modulated by light for the building of a more sensitive and broader photoreceptive system (Hannibal et al., 2005, Hannibal, 2006). It is therefore plausible that Tbr2^+^ ipRGC reservoir allows Opn4 expression to be dynamically regulated to cope with the need to adapt to the distinct lighting conditions in different geographic locations and/or seasons (Peirson et al., 2009). Our data that Tbr2 is essential for Opn4 expression supports this notion. Together, these data strongly support our conclusion that Tbr2^+^ RGCs are ipRGCs and that it would be of interest to study Tbr2’ roles in regulating the presumptive adaptive expression of *Opn4* in response to differential lighting conditions.

### Tbr2 expression marks all known ipRGC subtypes

Within Tbr2^+^ RGCs, we identified 2 types of OFF-layer stratified RGCs, several types of bistratified RGCs, and 3 types of ON-layer stratified RGCs. The predominant type of the OFF-layer Tbr2^+^ RGCs has been described previously as M1 ipRGCs, which have sparsely branching, monostratified dendritic ramifications in the outermost IPL and with variable dendritic field sizes and IMP profiles. This M1 type responds to a wide range of light intensity with unknow mechanisms. We found that M1-ipRGCs may be further divided into three subtypes: M1n, M1r and M1s. Among them, the M1s has a much lower input resistance than M1n and M1r. The exact relationship between IMP profiles, dendritic morphology, and dynamic light responses was not pursued further here but derserve future attention. The second and rare type of OFF-layer stratified RGCs was discovered in *Tbr2^CreERT2^:Pou4f1^CKO^* retinas. The co-expression of Tbr2 and Pou4f1 marks this unique RGC subtype with bushy and small dendritic arbors stratifying into the OFF sub-laminae slightly below the IPL/INL boundary. The overall morphology of this RGC subtype is reminiscent of that of types “d” and “j” Pou4f1^+^ RGCs (Badea and Nathans, 2011). We have also encountered a few of this RGC subtype in *Tbr2^CreERT2/+^: Rosa^iAP^* retinas, and it is likely that more of this type of RGC were buried in the dense dendritic AP signals from dACs. The intrinsic photosensitivity of this Tbr2^+^ RGC subtype is not determined, but it can be done by generating new mouse lines allowing intersectional fluorescent labeling. We encountered various types of bistratified Tbr2^+^ RGCs, which matched well with the published M3 and M6 types, Finally, we found 3 types of ON-layer stratified Tbr2^+^ RGCs that morphologically correlated well with known M2, M4, and M5 ipRGCs. This confirms that Tbr2^+^ RGCs are indeed ipRGCs. In agreement with our findings, a recent study using single-cell RNA sequencing (scRNA-seq) to profile RGC transcriptomes in P5 mouse retinas also showed that *Tbr2* expression is enriched in 7 (out of 40) RGC clusters (clusters 5, 6, 25, 26, 33, 37, 40) (Rheaume et al., 2018), while *Opn4* expression was expressed in 6 of the Tbr2^+^ clusters (higher levels in clusters 6, 25, and 33, and medium to low levels in clusters 5, 26, and 37). Furthermore, our birth-dating data suggest that the onset of ipRGC differentiation matches well with that of Tbr2^+^ RGCs (data not shown). Together, our data suggest that the terminal fate of ipRGC types is already determined at this early developmental stage, and thus Tbr2 is likely involved in ipRGC fate determination. Future research to compare the molecular profiles within each Tbr2^+^ cell type at different developmental stages will help elucidating the genetic network controlling the specification and differentiation of each of the ipRGC subtypes.

### Role of Tbr2 in regulating Opn4 expression

A well documented function of T-box transcription factors is to regulate molecular pathways involved in cell migration and cell-cell and cell-environment interactions (Strumpf et al., 2005, Russ et al., 2000, Costello et al., 2011, Nowotschin et al., 2013, Shen et al., 2014, Pearce et al., 2003, Ciruna and Rossant, 2001). For example, *Tbr1*, a paralogue of *Tbr2*, is necessary for neuronal migration, axon projection, establishment of regional and laminar identity in developing cortex and olfactory bulb, as well as RGC dendritic morphogenesis (Hevner et al., 2001, Huang et al., 2014, Kiyama et al., 2019, Liu et al., 2018). In mouse retinas, *Tbr2* expression appears in a few RGCs that have migrated to the innermost region of GCL at ~E14, suggesting that it plays a role in early steps of RGC development, such as cell determination and axon pathfinding. However, how Tbr2 functions to achieve this and what are its downstream genes remain unclear. This study shows that deleting *Tbr2* in mature retinas leads first to downregulation of *Opn4* expression in ipRGCs, followed by their death, thus suggesting that in additional to the aforementioned developmental role, *Tbr2* is also essential for maintaining *Opn4* expression and ipRGC survival in adult. We expect similar function played by Tbr2 during development because it is expressed in differentiating RGCs prior to the onset of *Opn4* expression. Because one of the features of ipRGC subtypes is that each subtype expresses Opn4 at different levels, the remarkable extent to which there is Tbr2 occupancy on many *T*-elements in the *Opn4* locus raises the possibility that these Tbr2-bound elements may be involved in regulating *Opn4* transcription output differentially in different ipRGC subtypes. In conjunction with our previous study in which we showed that *Tbr2* is a direct downstream target of Pou4f2 and Isl1 (Mao et al., 2008, Wu et al., 2015), we demonstrated that the Pou4f2/Isl1-Tbr2-Opn4 regulatory hierarchy is necessary for ipRGC formation. However, how the diverse ipRGC subtypes arise from this simple lineal gene regulatory hierarchy to give rise to distinct dendritic morphologies, retinofugal projections, intrinsic membrane properties, and molecular profiles remain unanswered. Nevertheless, the close association of Tbr2 with all ipRGC subtypes suggests a pivotal role it plays in this process.

In summary, we have demonstrated that Tbr2-expressing RGCs are morphologically indistinguishable from ipRGCs and that *Tbr2* is essential for maintaining *Opn4* expression and the survival of adult ipRGCs. Future studies should investigate Tbr2’s co-regulators and downstream effectors in controlling the formation and maintenance of all ipRGC subtypes. Our generation of a novel *Tbr2* reporter mouse line provides a unique opportunity to study several biological questions regarding RGC specification, maturation, and terminal differentiation. One example we have shown in this study is that a single M5-like Tbr2^+^ RGC sends collateral input to multiple brain targets in the IGL, vLGN, and OPN, suggesting a single Tbr2^+^ RGC may relay diverse photic information that requires different photo-response kinetics and threshold sensitivities, underlining the multiple roles of a single ipRGC in non-image forming vision. The comprehensive brain targeting information of a single RGC has been reported for M1-ipRGCs (Fernandez et al., 2016), but such knowledge is still lacking for other ipRGC subtypes. It will be of interest to trace retinofugal projections of all Tbr2^+^ RGC subtypes at the single cell level to fill in the knowledge gap in the anatomical architecture and the underlying functional implications for different ipRGC subtypes.

## AUTHOR CONTRIBUTIONS

CAM, CKC, CW, TCB and SCM designed experiments. CAM, CKC, CW, TK, NW, PP, and SCM executed experiments. CAM designed and generated Tbr2^TauGFP-IRESCreERT2^ mice. TCB provided mice and reagents. CAM and CKC wrote the paper.

## Acknowledgments

This work was supported by grants from the National Institutes of Health-National Eye Institute to C.A.M. (EY024376) and C.K.C (EY026930), C.M.W. (EY021958), S.C.M. (EY006515), and in part by National Eye Institute Vision Core Grants P30EY028102 (UTHealth) and P30EY002520 (BCM). The Ophthalmology Department of Baylor College Medicine received an unrestricted grant from Research to Prevent Blindness, Inc. CKC is the Alice McPherson Retinal Research Foundation Chair at BCM. We acknowledge Dr. Jan Parker-Thornburg and Genetically Engineered Mouse Facility at The University of Texas MD Anderson Cancer Center in making *Tbr2^TauGFP-IRESCreERT2/+^* mouse line, Dr. Samer Hattar at NIMH for sharing *Opn4^Cre^* mouse, Dr. King-Wai Yau at Johns Hopkins University for sharing *Opn4^TauLacZ^* mouse, and Dr. Kimberly Mankiewicz at UTHealth for proofreading the manuscript.

